# Charge neutralization of the active site glutamates does not limit substrate binding and transport by EmrE

**DOI:** 10.1101/2022.08.17.503475

**Authors:** Peyton J. Spreacker, Merissa Brousseau, Grant S. Hisao, Mohammad Soltani, James H. Davis, Katherine A. Henzler-Wildman

## Abstract

EmrE, a small multidrug resistance (SMR) transporter from *E. coli*, confers broad-spectrum resistance to polyaromatic cations and quaternary ammonium compounds. Previous transport assays demonstrate that EmrE transports a +1 and a +2 substrate with the same stoichiometry of 2 protons:1 cationic substrate. This suggests that EmrE substrate binding capacity is limited to neutralization of the two essential glutamates, E14_A_ and E14_B_ (one from each subunit in the antiparallel homodimer), in the primary binding site. Here we explicitly test this hypothesis, since EmrE has repeatedly broken expectations for membrane protein structure and transport mechanism. We previously showed that EmrE can bind a +1 cationic substrate and proton simultaneously, with cationic substrate strongly associated with one E14 residue while the other remains accessible to bind and transport a proton. Here we demonstrate that EmrE can bind a +2 cation substrate and a proton simultaneously using NMR pH titrations of EmrE saturated with divalent substrates, for a net +1 charge in the transport pore. Further, we find that EmrE can alternate access and transport a +2 substrate and proton at the same time. Together, these results lead us to conclude that E14 charge neutralization does not limit the binding and transport capacity of EmrE.

## Introduction

Determining the substrate profile of a transporter is important for understanding which molecules the transporter can or cannot move across the membrane. This task is more challenging in the case of promiscuous transporters, such as the multidrug resistance efflux pumps, where substrates span a very broad chemical space. In the case of ion-coupled transporters, binding of the small molecule substrate must also trigger the coupled conformational changes necessary for alternating access to allow transport to occur (Fig. 1A). Here we explore the importance of substrate charge on ligand recognition, molecular gating, and transport for the small multidrug resistance transporter EmrE. This transporter has become a model system for studying multidrug recognition and the mechanism of secondary active transport.

**Figure 1:**
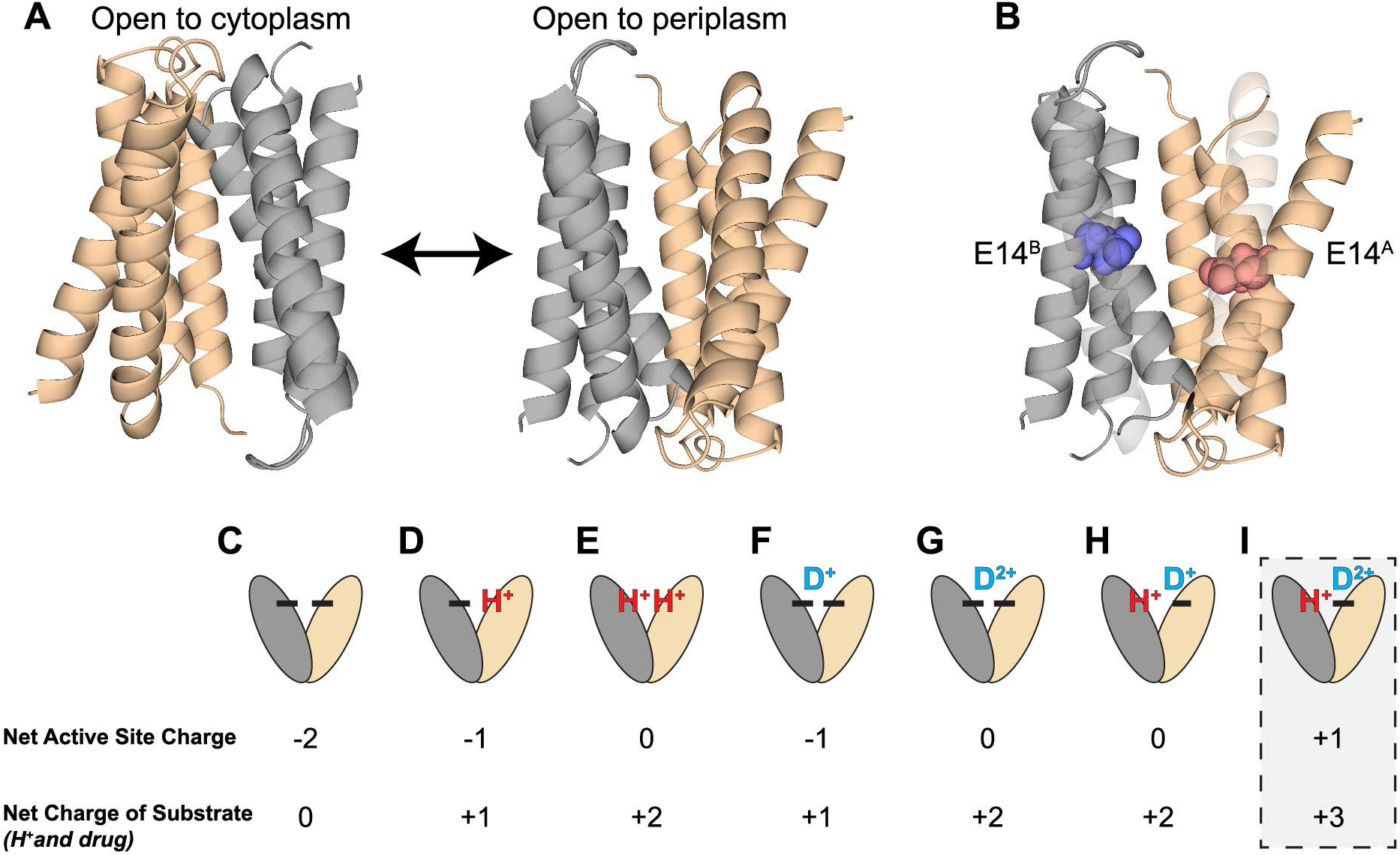
E14 defines the proton and drug binding site in EmrE. The two subunits in the antiparallel asymmetric EmrE homodimer (chain A, tan; chain B, gray) swap conformations to alternate access of the transport pore between open-in and open-out (A). Chain designations are based on PDB 3B5D and the structure is a molecular dynamics refinement of the structure in explicit lipid (30). Two glutamate residues, E14^A^ and E14^B^, one on each chain of the antiparallel homodimer make up the binding site of EmrE (B). The various potential states of EmrE (C-I) are depicted along with their corresponding net active site charge and net change of bound substrates based on biophysical and NMR measurements of EmrE interaction with the +1 substrate TPP^+^. The putative state of EmrE simultaneously bound to a +2 drug substrate and a proton (G) is the focus of this work and would result in a net positive charge within the transport pore.

The primary binding site for small molecule substrates and protons in the EmrE transporter is centered on two essential glutamates, one from each subunit in the antiparallel homodimer, E14_A_ and E14_B_ (Fig. 1B) (1–3). Glutamate residues are among the most common amino acids found in the active sites of enzymes and transporters where they may hydrogen bond or interact electrostatically with the substrate. Mutation of E14 to anything other than aspartate abolishes binding of protons, binding of the known antiported substrates, and the ability of EmrE to transport and confer resistance to those substrates (1, 2). EmrE can bind and transport both +1 and +2 cations (4). Radiolabeled transport assays measured electrogenic transport of a +1 substrate (tetraphenylphosphonium, TPP^+^) and electroneutral transport of a +2 substrate (methyl viologen, MV^2+^). The data for these two substrates is consistent with EmrE acting as a tightly coupled antiporter with a consistent 2 H^+^ to 1 drug stoichiometry for both +1 and +2 drug substrates (4). Tightly coupled antiport with a single binding site is achieved simply by invoking a model of strict competition between proton and small molecule substrates for binding to that single site and assuming alternating access only occurs when a substrate (small molecule or proton) is bound. In the case of EmrE, binding of a positively charged small molecule substrate or proton(s) to the primary binding site defined by E14 will reduce the net charge in the core of the transporter transmembrane region, suggesting a natural mechanism by which substrate binding lowers the energy barrier for alternating access (Fig. 1).

However, more recent NMR studies, biophysical experiments and kinetic simulations revealed that EmrE is a loosely coupled transporter capable of binding a +1 small molecule substrate and proton simultaneously and alternating access with +1 substrate and proton bound (5, 6). With two E14 residues in the asymmetric homodimer, Fully deprotonated EmrE has a net active site charge of −2 (Fig. 1C), with successive protonation (7), leading to net active site charges of −1 or 0 (Fig. 1D, 1E). Binding of a +1 drug-substrate occurs asymmetrically with the substrate more closely associated with one subunit (5) and results in a net active site charge of −1 (Fig. 1F). Simultaneous proton binding to the opposite glutamate results in a net active site charge of 0 and is still consistent with a model where charge neutralization of the glutamate sidechains enables alternating access and transport.

Binding a +2 drug-substrate alone (Fig. 1G) leads to a neutral active site (4). Here, we use NMR to test whether EmrE can bind a +2 drug-substrate and a proton at the same time (Fig. 1I) and assess whether alternating access occurs when there is a net positive charge in the transport pore, as would be needed to transport of a +2-drug substrate and proton across the membrane simultaneously. These experiments will determine whether EmrE can perform loosely coupled transport of +2 small molecule substrates, or loose coupling is only possible in the case of +1 substrates. It will also provide insight into whether net charge in the transport pore is a key factor in lowering the energy barrier for alternating access and enabling transport to occur.

## Results

To test the hypothesis that neutralization of the two active site glutamates, E14A and E14^B^, limits the binding capacity of EmrE within the transport pore, we performed three types of experiments. First, we used NMR pH titrations to directly assess whether EmrE can bind both a +2 small molecule substrate and a proton simultaneously, resulting a net +1 charge in within the hydrophobic transport pore. Second, using the results of the NMR pH titrations to determine appropriate conditions for simultaneous binding of +2 small molecule substrate and proton, we assessed whether EmrE could alternate access with a +1 net charge in the transport pore using ZZ-exchange NMR dynamics experiments. Third, we used SSME to assess whether we could detect transport indicative of EmrE simultaneously moving a +2 drug-substrate and proton across the liposomal membrane.

### EmrE binds propidium and proton simultaneously

We first tested whether EmrE can simultaneously and proton and a well-established +2 substrate, propidium (PP^2+^). We performed NMR-monitored pH titrations of PP^2+^-saturated EmrE, the same approach previously used to demonstrate simultaneous binding of +1 small molecule substrate and a proton (5). The pH titration spanned from pH 4.5 to 8.0 (Fig. 2, Fig. S1) and the spectra show chemical shift perturbations indicative of a protonation event occurring within this pH range. Chemical shift perturbations occur due to the change in electrostatic environment upon proton binding but can also occur due to binding PP^2+^ or coupled conformational changes. We therefore performed a series of control experiments. Drug-like substrates have lower affinity for EmrE at low pH, so it is important to verify that PP^2+^ remains bound at low pH and the chemical shift perturbations observed during the pH titration are due to protonation/deprotonation. At pH 5.0 (Fig. 2A, red), the PP^2+^-bound spectrum shows residual chemical shift perturbations (CSPs) when compared to the drug-free spectrum at this pH (Fig. 2A, black), confirming that EmrE binds PP^2+^ and H^+^ simultaneously and does not release the small molecule substrate upon proton binding. At high pH (Fig. 2B), CSPs are also present between drug-free (black) and PP^2+^-bound (blue) WT-EmrE confirming direct binding of propidium as previously shown (8). It should be noted that if the 2+ charge could neutralize the E14 pair in the active site, then no pH-dependent chemical shift perturbations should be observed when EmrE is saturated with PP^2+^.

**Figure 2:**
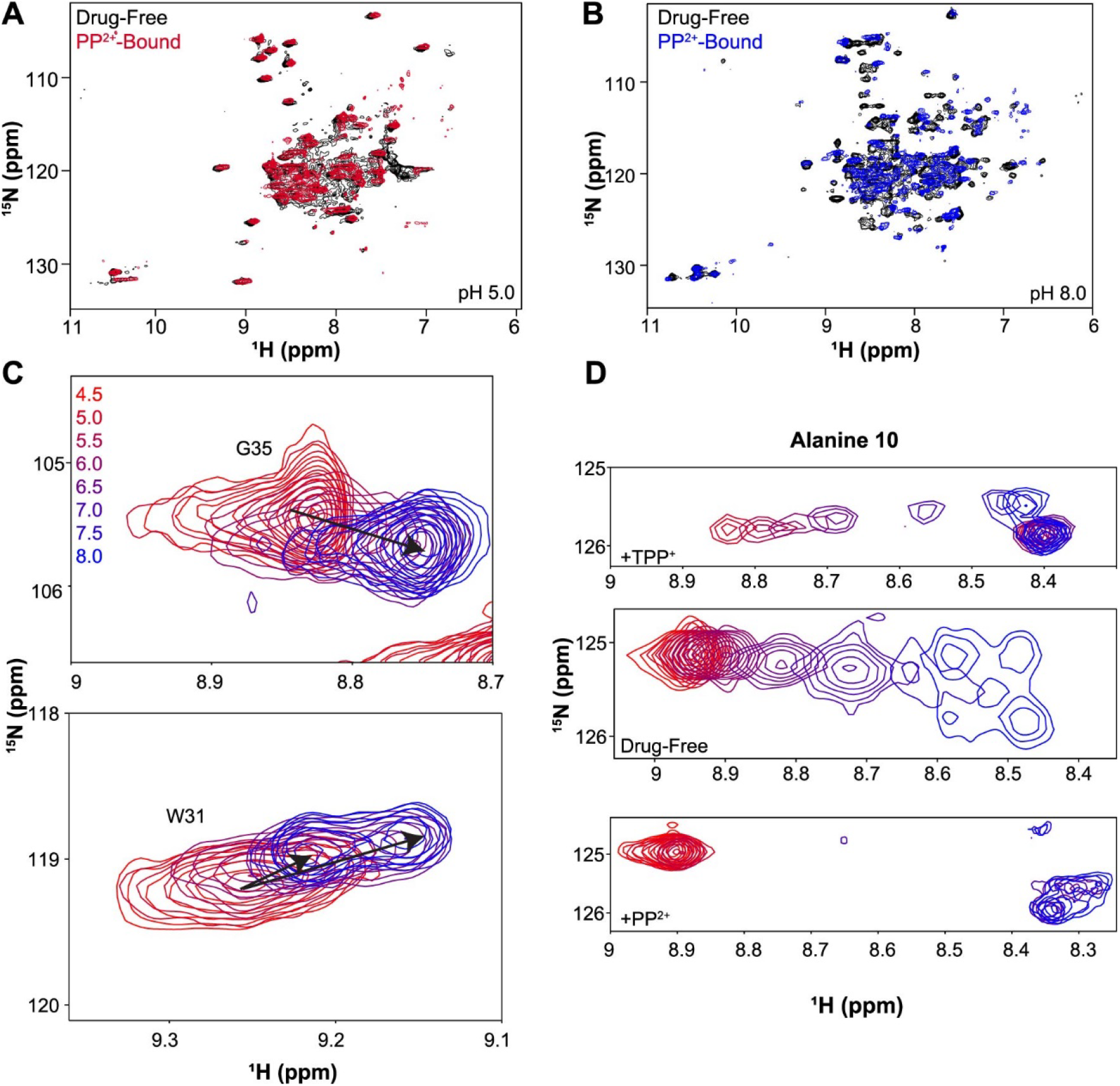
EmrE binds propidium^+2^ and proton simultaneously. ^1^H-^15^N TROSY monitored pH titration of EmrE saturated with PP^2+^ at 45°C reveals that EmrE can bind proton and PP^2+^ simultaneously. Spectra were acquired on an 800 MHz NMR spectrometer with EmrE solubilized in q=0.33 DMPC/DHPC isotropic bicelles. Chemical shift changes occur as a function of pH, indicative of protonation at E14 in the PP^2+^-bound state. Chemical shift differences between substrate-free and PP^2+^-saturated EmrE at low pH (pH 5.0, A) and high pH (pH 8.0, B), confirm that EmrE binds PP^2+^ in both the protonated and deprotonated form and that proton binding at low pH does not prevent PP^2+^ binding. Chemical shift perturbations of A10 and other residues as a function of pH highlight the impact of E14 protonation colored by the corresponding pH condition (C, D). Alanine 10 (A10) is close in space to E14 and removed from other protonatable residues (D). The full pH titration spectrum can be found in Figure S1.

An additional control experiment was performed to confirm that the observed CSPs reflected protonation and PP^2+^ binding at E14. Mutation of this critical residue in E14Q-EmrE eliminates most of the pH-depending chemical shift changes, with only the C-terminal H110 and nearby residues titrating within the near-neutral pH range (Fig. S3). This demonstrates that the widespread pH- and PP^2+^-dependent CSPs observed upon pH titration of PP^2+^-saturated EmrE or comparison of apo and PP^2+^-saturated EmrE at low- and high-pH reflect proton and PP^2+^ binding at E14. Residues near E14, such as G9, G17, and S43 (Fig. S1A) undergo large shifts or split during the pH titration of PP^2+^-bound WT-EmrE. These residues are important in substrate binding and specificity (9) and have previously been shown by our lab to be perturbed upon the binding of another small molecule substrates, TPP^+^ (5, 7). We have previously shown that these residues (7) directly reflect protonation events at E14 and not long-range conformational changes that may occur upon protonation.

While multiple peaks in proximity to E14 display similar behavior, as can be seen in careful analysis of the full spectra (Fig. S1), we illustrate the results by focusing on alanine 10 (A10). A10 is an ideal residue to track the impact of E14 protonation by NMR because it is well-resolved in the spectrum and close in space to E14, only one turn away on the same side of transmembrane helix 1. Comparing the A10 pH-dependent chemical shifts of drug-free, TPP^+^-bound, and PP^2+^-bound WT-EmrE (Fig. 2D) reveals a clearer picture of how protonation affects the rate of alternating access. EmrE functions as an asymmetric antiparallel homodimer, and distinct chemical shifts are observed for each subunit of the homodimer because they have different structures. The two subunits swap conformations, resulting in a homodimer that has the same overall structure but is open to the opposite side of the membrane (Fig. 1B) (10). This alternating access process can be easily observed by NMR and is pH-dependent. As previously described (7), alternating access is fast (≥200 s^-1^) in the fully protonated state (no drug substrates, low pH), resulting in a single peak at the average chemical shift Fig. 2D, middle). As the pH increases, the protein enters intermediate and then slow-intermediate exchange, as observed by the line broadening and then splitting into separate two peaks reflecting the two distinct subunit conformations (marked A and B), with exchange peaks (marked x) reflecting the fact that this is not fully in the slow exchange limit, with alternating access occurring at a rate of ≈20-40 s^-1^.

We previously studied the simultaneous binding of EmrE with the +1 substrate, TPP^+^, and H^+^ (5). We found that TPP^+^ interacts with EmrE asymmetrically (Fig. 2D, top) to subunit A, while subunit B remains accessible to protonation with pH-dependent CSP of nearby residues revealing a pKa of 6.8 ± 0.1. With TPP^+^-bound, alternating access remains in the slow-exchange regime throughout the tested pH range. The +2-substrate PP^2+^ has a much lower binding affinity than TPP^+^ (8) and PP^2+^-bound EmrE has significant line broadening of residues in the transport pore. This line broadening is likely due to the motion of the planar PP^2+^ causing fluctuating ring currents. TPP^+^ is highly dynamic even when bound tightly and asymmetrically within the transport pore (5, 11, 12), but the symmetric nature of this substrate limits the impact of this motion on the observed NMR resonances. Despite these challenges, pH-dependent CSPs of PP^2+^-bound EmrE (Fig. 1D, bottom) demonstrate a completely different behavior from previous experiments. Alternating access is in the fast exchange limit at low pH and in slow-intermediate exchange at high pH (Fig. 1D, Fig. S2A, B), most similar to drug-free EmrE (Fig. 1A). At intermediate pH, the spectra are more severely broadened than for drug-free EmrE, as would be expected due to the additional factor of PP^2+^ motion within the transport pore in the vicinity of A10. Inspection of the H110 backbone amide peak (Fig. S1, B) shows a smooth titration and distinct pKa, confirming that the behavior of A10 reflects the protonation of E14 and not the water-exposed C-terminal H110 residue. The slow-intermediate timescales observed at neutral pH make the determination of the PP^2+^-bound pKa impossible, and it is not clear whether proton and PP^2+^ bind asymmetrically to either subunit of the homodimer (as observed for TPP^+^), but protonation does occur in the PP2+-bound state.

### EmrE has similar binding affinities and exchange rates for dTPP^+^ and dTPP^2+^

While TPP^+^ and PP^2+^ are well-known substrates of EmrE, using these substrates to assess the impact of substrate charge is complicated because these small molecules have vastly different molecular weights and chemical structures. We have previously shown that small molecule size, charge and hydrophobicity all affect binding affinity, rate of alternating access and transport by EmrE (8). To focus more cleanly on the impact of substrate charge on the ability of EmrE to simultaneously bind proton and a polyaromatic cation substrate, we synthesized two compounds designed to minimize the difference in size, shape, mass, and hydrophobicity (Fig. 3). Solution NMR spectroscopy was then performed on these compounds to confirm their structures (Fig. S4, Fig. S5). To properly quantify these ligands in solution, extinction coefficients of 1,4-phenylenebis (triphenylphosphonium) bromide (dTPP^2+^) and triphenyl[4-(triphenylmethyl)phenyl]phosphonium iodide (dTPP^+^) were determined using UV-Vis spectroscopy at 269 nm and 276 nm. The traces of the UV-Vis spectra can be found in Fig. S6 and Fig. S7, and a summary of each set of measurements is in Table S1.

**Figure 3:**
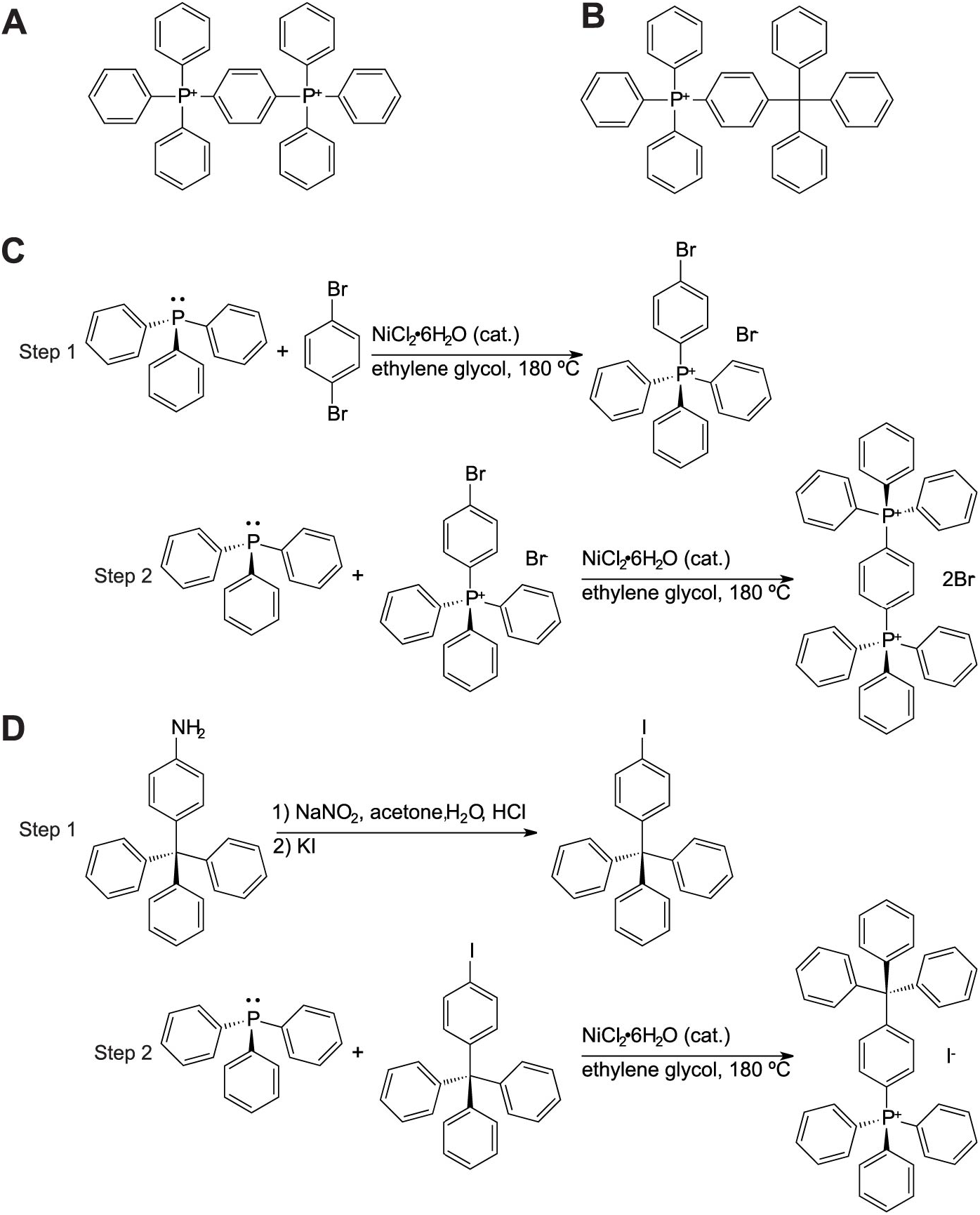
Synthesis of dTPP^+^ and dTPP^2+^. (A) dTPP^2+^ has a triphenyl phosphonium group off the para position on one of the TPP^+^ phenyl rings, resulting in an asymmetric molecule with two phosphorus centers and a net 2+ charge. (B) dTPP^+^ is similar in structure to dTPP^2+^ but has a carbon atom instead of a phosphorus atom in one of the centers, which results in a charge of 1+. Panels C and D show the synthetic schemes for dTPP^2+^ and dTPP^+^, respectively.

**Figure 4:**
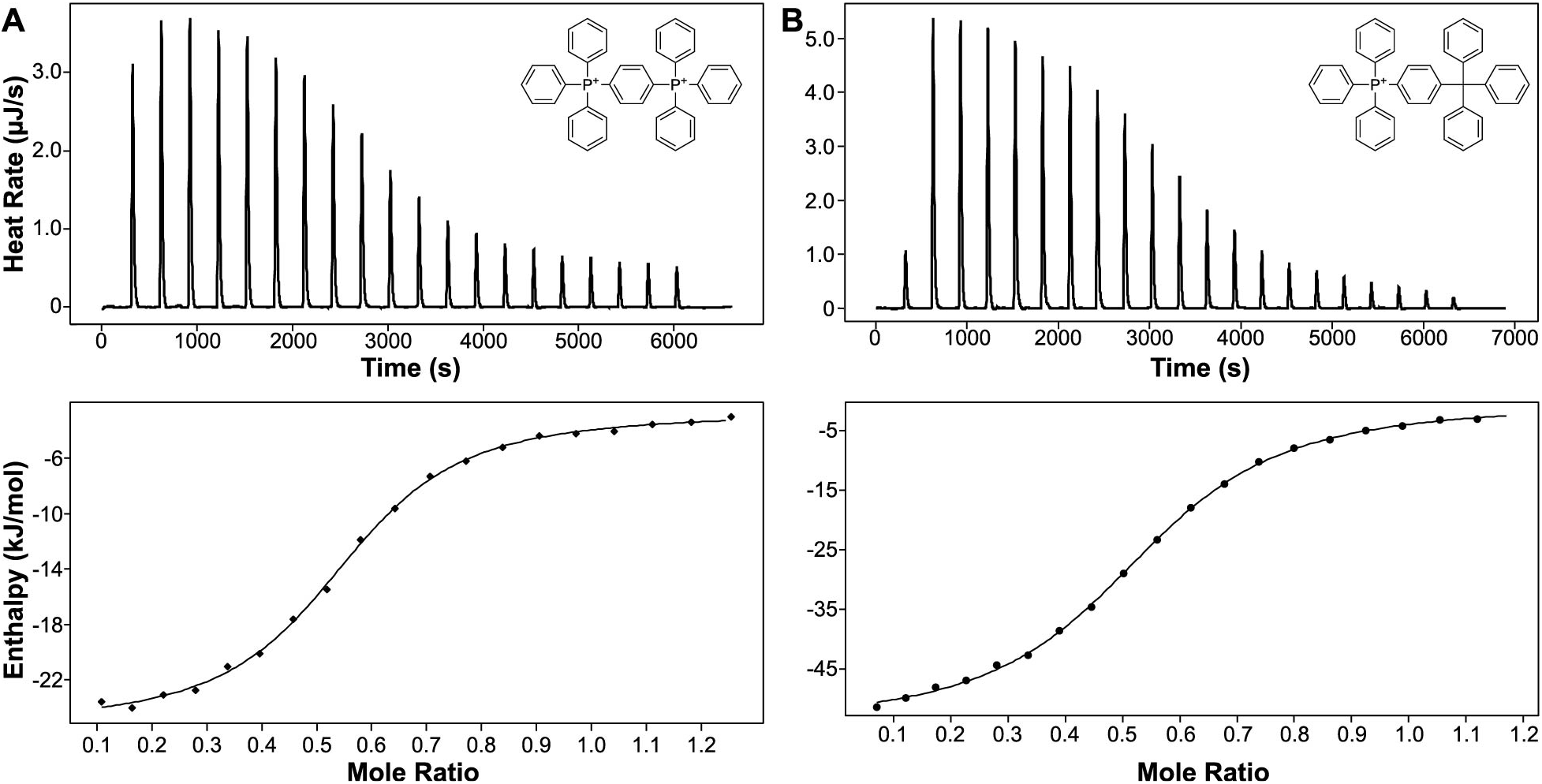
EmrE binds dTPP derivatives with similar affinity. Representative isothermal titration calorimetry and binding curves for dTPP2+ (A) and dTPP+ (B) binding to EmrE in isotropic bicelles. The dissociation constants (*K*_d_), enthalpy (ΔH), and entropy (ΔS) of binding were determined at pH 7.0 and 45 °C. Each measurement was done in triplicate.

To first demonstrate that EmrE can bind to these substrates, isothermal titration calorimetry (ITC) experiments were performed in isotropic bicelles for each ligand (dTPP^2+^, dTPP^+^, ^2^H-dTPP^2+^). The binding constants (K_D_) of dTPP^2+^, dTPP^+^, and ^2^H-dTPP^2+^ were determined at pH 7.0, 45 °C to be 8.1 ± 0.4 μM, 9.3 ± 0.3 μM, and 13 ± 1 μM respectively (Fig. 3). Table II summarizes the average dissociation constants, enthalpy, entropy, Gibbs free energy, and stoichiometry of EmrE binding to each respective ligand. The values for each experiment as well as the traces for each experiment can be found in Fig. S8 and Table S2. The determined K_D_ values confirm that each ligand binds EmrE with a relatively similar affinity. Though small differences in the K_D_ are observed for each ligand, the values remain within the same order of magnitude and are thus considered minimal. This is especially true in the context of known EmrE ligands, for which K_D_ values can range from as low as 40 nM for strong binders and as high as 130 μM for weak binders (8).

### EmrE can bind dTPP^+^ or dTPP^2+^ and proton simultaneously

We first performed a pH titration of dTPP^+^-bound EmrE from pH 4.5 to 8.0 to observe the effect of pH on EmrE bound to the +1 derivative (Fig. 5). Comparing the shifts observed throughout the protein (SI Fig. 9) to previously acquired chemical shifts of E14Q- and E14D-EmrE (3, 7), we can separate the effects of protonation at H110 from protonation at E14. There are additional shifts that occur in the presence of dTPP^+^ that are not in the E14Q-EmrE spectra (Fig. S3). As described above, this is indicative of simultaneous proton and drug binding at the primary E14 site within the transport pore of EmrE. As illustrated by the A10 resonances (Fig. 5A), the pKa of dTPP^+^-bound EmrE is higher than for TPP^+^-bound EmrE, with titration not completed at pH 7.5-8.0. The alternating access dynamics of the protein are in intermediate-fast exchange at low pH, as observed for drug-free and PP^2+^-bound EmrE, and many residues in the active site have additional line broadening. As for PP^2+^, this is expected given what is known about the motion of the substrate within the transport pore and the less symmetric structure of dTPP^+^ compared to TPP^+^. The resulting line broadening prevents more detailed analysis at low pH, but binding is evident when comparing WT-EmrE without (Fig. 5B, black) or with dTPP^+^ (Fig. 5B, red) at pH 5.0.

**Figure 5:**
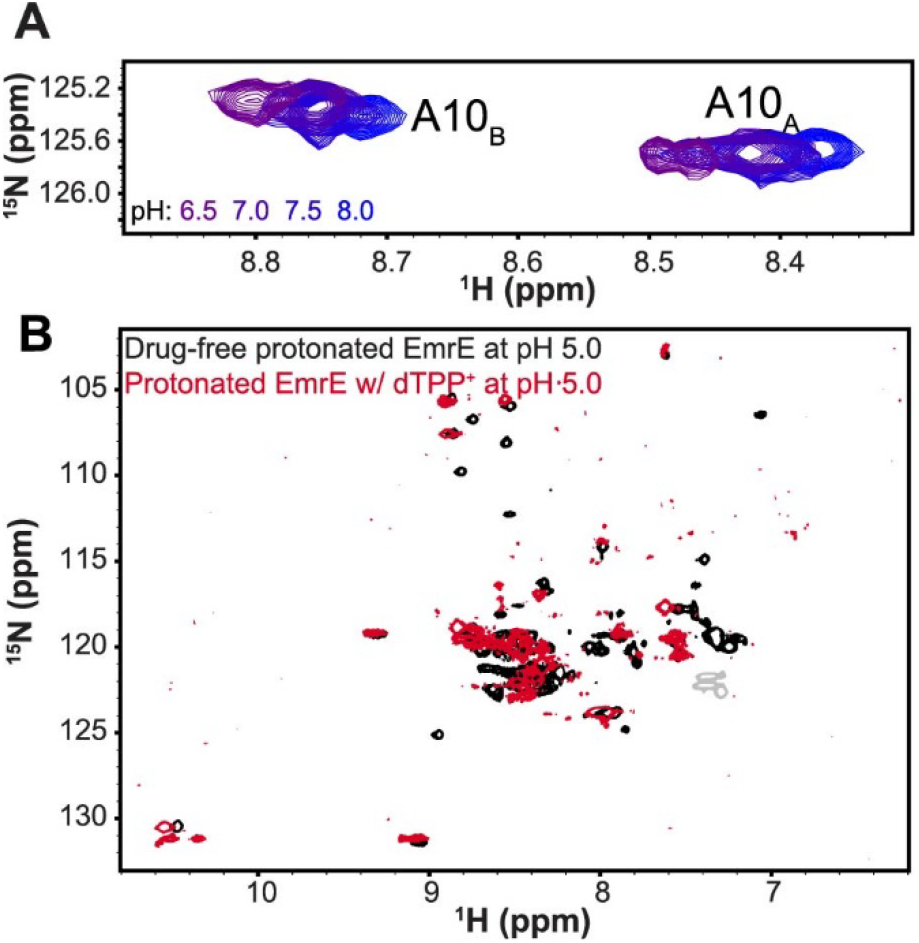
EmrE binds dTPP^+^ with increased dynamics at low pH. (A) ^1^H-^15^N TROSY spectra of EmrE saturated with dTPP^+^ as a function of pH show chemical shift perturbations indicative of proton binding to dTPP^+^-saturated EmrE. (B) pH titration of A10 resonances at high pH are similar to those observed for pH titration of EmrE bound to TPP^+^ (ref) or PP^2+^ (Fig. 2). (C) While there is increased line broadening at lower pH conditions, dTPP^+^ still binds to EmrE as shown by chemical shift differences between the bound (red) and apo (black) EmrE spectra at pH 5.0.

To characterize the pH dependence of EmrE with the +2 charged ligand species, dTPP^2+^, another NMR pH titration was conducted (Fig. 6). This substrate has a lower affinity compared to TPP^+^, so higher concentrations were required for saturation and ^2^H-dTPP^2+^ was used to minimize substrate peaks in the proton-detected NMR spectra. Fig. 6A shows examples of chemical shift perturbations observed in the spectra for two residues in spatial proximity to E14 (residues A10 and V66). The full overlay of the titration spectra can be found in SI Fig. 10. The ^1^H and ^15^N chemical shifts of 7 residues were fit globally against pH (Fig. 6B), allowing us to determine a single pKa of EmrE bound to dTPP^2+^ of 6.2 ± 0.1. As observed for the +1 substrate TPP^+^, alternating access is in slow exchange throughout the pH range and substrate binding is asymmetric, with dTPP^2+^ more closely associated with E14 in one subunit (A), protecting that subunit from protonation-dependent chemical shift changes while E14 in the other subunit (B) titrates with pH. The differences in pKa values, (6.2 ± 0.1 for dTPP^2+^, 6.8 ± 0.1 for TPP^+^ (5)), is evidence of how substrate size and charge shift the equilibrium of proton binding. The pH titrations of EmrE bound to either +2 substrate (PP^2+^ or dTPP^2+^) show clear evidence of simultaneous proton binding and contradict the hypothesis that charge neutralization of the active site E14 residues limits substrate binding to EmrE.

**Figure 6:**
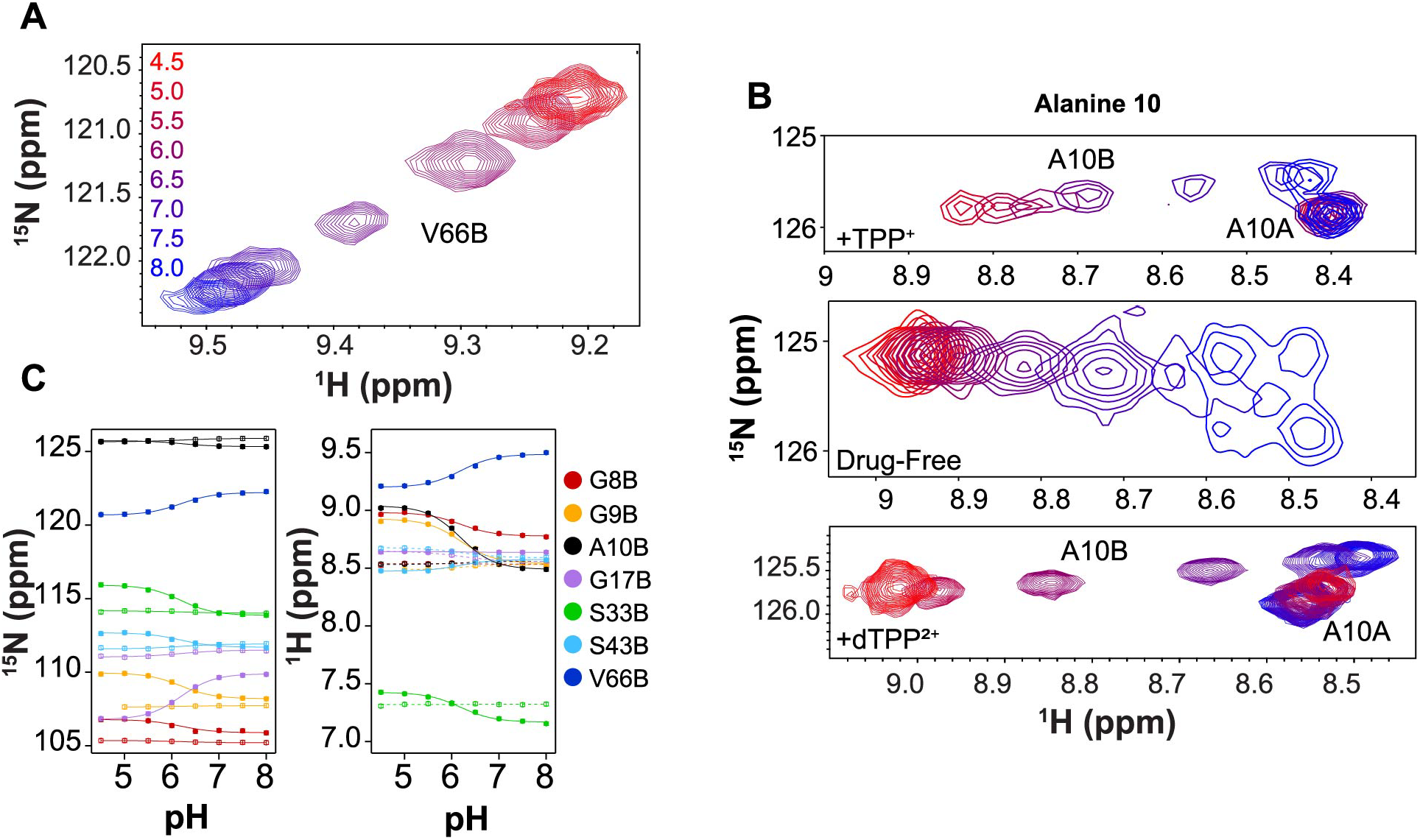
EmrE binds dTPP^2+^ and proton simultaneously. (A) ^1^H-^15^N TROSY spectra of EmrE saturated with dTPP^2+^ as a function of pH show chemical shift perturbations indicative of proton binding to dTPP^2+^-saturated EmrE. (B) Highlight of pH titration for the A10 resonances. (C) Chemical shift perturbations (CSPs) for seven residues were simultaneously fit to determine the single pKa value of 6.2 ± 0.1.

### EmrE alternates access with dTPP^2+^ and proton bound simultaneously

As with +1 substrates, simultaneous binding of a +2 substrate and proton only impacts EmrE activity if the transporter can alternate access to simultaneously move the substrates across the membrane. To investigate the ability of EmrE to transport a +2 drug substrate and a proton at the same time, we first needed to test whether EmrE could alternate access with dTPP^2+^ and proton bound to the active site. We therefore performed methyl ZZ-exchange spectroscopy on dTPP^2+^-saturated EmrE at pH 5.0 to ensure simultaneous proton binding (Fig. S11). At low pH, an exchange rate of 6.1 ± 0.5 s^-1^ was measured for dTPP^2+^-bound WT-EmrE (Fig. S11A). At high pH, dTPP^2+^- and dTPP^+^-bound EmrE had alternating access rates of 3.1 ± 0.1 s^-1^ and 3.5 ± 0.1 s^-1^, respectively (S11B, C). Together, these results demonstrate that indeed EmrE can alternate access with a +2 drug substrate and proton bound to its active site.

### Ethidium is transported in a coupled antiport mechanism by EmrE

To examine the effect of charge on EmrE-mediated transport, we sought to characterize the transport behavior of well-characterized +1 and +2 substrates using solid-supported membrane-electrophysiology (SSME). For the +1 substrate, we chose the common low affinity EmrE ligand, ethidium (Eth^+^). Prior efforts to screen for EmrE-transported substrates with SSME introduced drug in the absence of a pH gradient, relying on the drug gradient across the membrane to drive net charge movement according to the dominant transport mode (13). Since Eth^+^ contains a single positive charge and is transported by coupled antiport with a net 2:1 proton:drug stoichiometry, this results in a detectable electrogenic transport opposite the driving drug gradient.

Eth^+^ was excluded from analysis in the prior work because of anomalous signals in transport experiments with empty liposomes, likely due to association of ethidium with the lipid head groups under the high drug concentrations used in that work. Here, we examined Eth^+^ transport specifically by maintaining drug concentrations well below the expected binding constant, minimizing signal from potential weak interactions with empty liposomes. Even in the absence of any proton gradient, the introduction of the drug to the outside of the liposome creates a gradient of sufficient magnitude to initiate transport, with two protons moving out of the liposome for every Eth^+^ molecule transported into the liposome (Figure 6A). This is seen as the net movement of one positive charge per transport cycle moving out of the liposome, resulting in a negative current. Trials for the different WT-EmrE and empty liposome-containing sensors confirm that the observed signal under these conditions is transport dependent on the protein as empty controls show minimal signal (Figure S12).

Having established that Eth^+^ transport can be observed by SSME, we next sought to confirm that antiport is the predominant transport mode. Examining the reversal potentials of coupled substrates has become a standard way to characterize the direct stoichiometry of coupling for electrogenic transport (14, 15). However, a more general manipulation of the gradients of coupled substrates can provide a qualitative means of examining the dominant transport mode for a substrate. If a constant, inward-facing proton gradient is maintained, alterations in the magnitude and direction of the drug gradient for a given substrate can reveal the most favorable transport mode for that substrate based on the different magnitude and direction of flux expected under each condition for antiport, symport or uncoupled uniport (16).

Assuming antiport is the dominant transport mode for Eth^+^, then the maximal transported charge would be observed when the proton and drug gradients are opposite to one another so that both favor antiport. With the same two-fold proton gradient under all conditions, reversing the much larger drug gradient should reverse the direction of net transport, but the total signal should be smaller because the two gradients are attempting to drive antiport in opposite directions. As expected, this is exactly what was seen for Eth^+^ with a 16-fold congruent drug gradient capable of reversing the direction of transport in the presence of a 2-fold proton gradient (Figure 6B) confirming antiport as the dominant transport mode for this substrate. While this analysis is incredibly useful for determining the transport mode of a substrate, the resulting signal is dependent on the observed transport being electrogenic. In the case of a +2 substrate, 2:1 antiport would be electroneutral and this approach cannot be used to demonstrate transport.

### SSME of methyl viologen supports simultaneous +2 drug/proton transport

We first assessed EmrE transport of methyl viologen (MV^2+^), a +2 substrate shown to be antiported by EmrE using radio-labeled transport assays (4). Performing standard transport experiments with an inwardly-directed MV^2+^ gradient and no proton gradient resulted in the expected lack of signal for 2:1 proton:drug antiport (Figure S13A). To characterize the electroneutral transport of MV^2+^, we exploited the promiscuous nature of EmrE transport made evident by the Free Exchange Model (5, 6). By generating gradients with a large enough magnitude, we can drive transport modes that may not be the most favorable, including proton uniport, and proton-drug symport (Figure 7A). While strict antiport of a +2 substrate for 2 protons results in a null signal, (black line), a strong inward-directed proton gradient paired with an inward-facing drug gradient of sufficient magnitude should be able to drive proton/drug symport, resulting in net movement of +3 inward (blue line). If the gradients are flipped such that the higher concentrations of proton and drug are inside the liposome, the signal will be of the same magnitude, but in the opposite direction (red line). While the NMR experiments discussed above support the ability of EmrE to bind a +2 substrate and proton simultaneously and even alternate access, we wanted to assess if this extends to simultaneous *transport* of proton and +2 substrate across a membrane. By comparing the magnitude of the signals for proton/drug symport (+3) and proton uniport (+2) for comparable gradients, we can begin to examine if the symport of a +2 substrate is, in fact, feasible as predicted by the free exchange model and NMR data.

**Figure 7:**
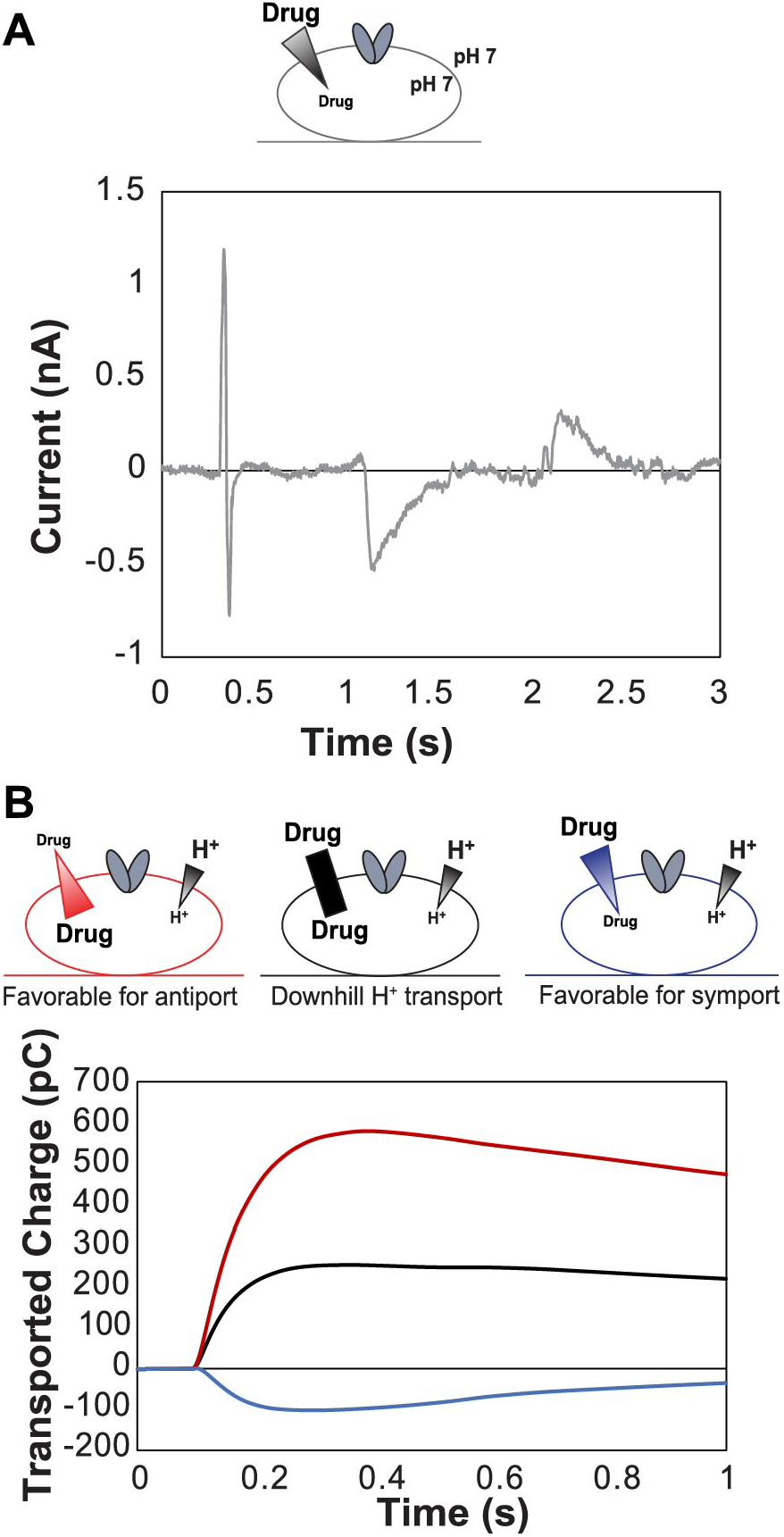
SSME enables characterization of electrogenic transport. Transport can be observed for transporters with moderate flux by adsorbing proteoliposomes onto a gold sensor and observing the combined current upon perfusion of various buffer conditions. With only a drug gradient present across the membrane, this gradient will drive transport according to the dominant mode. (A) For Eth^+^, this results in net antiport of approximately two protons for every Eth^+^ molecule resulting in a negative signal reflecting net transport of +1 out of the liposome per turnover. (B) The transport mode can be confirmed by performing experiments with disparate gradients such that each of the possible transport modes for EmrE (antiport, symport, and drug or proton uniport) will result in a specific pattern of transported charge.

Using the same drug concentrations that yielded no net charge movement under the antiport assay conditions, we tested whether we could detect symport when the proton gradient was aligned with the drug gradient and observed small signals indicative of net charge movement under these conditions (Figure S13B). This also confirmed that the proteoliposomes on the sensor were transport-competent and the null results in Fig. SI13A reflect electroneutral antiport. Empty liposomes were subjected to the same gradient conditions confirming that the small transport signals observed in Fig. SI13B are not due to association of the substrate with the liposome, since there is minimal current with empty liposomes (Figure S13C,D).

Since the free exchange model predicts that EmrE should be capable of performing proton uniport, we first tested whether we could drive uncoupled proton transport using a 10-fold proton gradient (pH 6 vs. pH 7) in the absence of any small molecule substrates. We observed current reflecting net charge movement down the proton gradient in liposomes containing WT EmrE but no significant current in empty liposomes (Fig. SI 14B), confirming that uncoupled proton transport is mediated by EmrE in the presence of a 10-fold proton gradient. Since EmrE is an antiparallel homodimer with one subunit oriented each way in the membrane, the preferred orientation in the membrane is not relevant. As expected, based on this unique structure, EmrE-mediated proton leak is identical but opposite when the direction of the 10-fold proton gradient is reversed (Figure 7B, top, light colors). These experiments were performed using the same sensors with extensive washes to change the internal buffer, and the identical magnitude of net charge transport confirms that this procedure is sufficient to fully exchange the internal buffer composition.

We next tested the impact of adding a drug gradient to test whether we could drive symport of +2 substrate and proton. Liposome internal buffer was exchanged to high pH (low proton concentration) and low drug concentration (pH 7, 250 nM MV^2+^). A buffer with low pH (high proton concentration) and high drug concentration (pH 6, 10 uM MV^2+^) was then perfused to create inward-directed 10-fold proton and 40-fold drug gradients (Figure 7B, top, dark blue) and the net charge movement was recorded. Finally, the internal buffer was exchanged to the higher concentration of drug and proton (pH 6, 10 uM MV^2+^) and the lower concentration buffer (pH 7, 250 nM MV^2+^) was perfused over the outside while transport was recorded (Figure 7B, top, dark red).

### SSME of propidium supports simultaneous +2 drug/proton transport

To ensure that these results are not specific only to MV^2+^, we tested a second +2 substrate, PP^2+^. PP^2+^ is more structurally similar to Eth^+^, and thus provides a better comparison of the impact of charge on substrate transport. For quantitative comparison of SSME data it is necessary to directly compare conditions on the same sensors because the loading of proteoliposomes onto the surface sensor surface is variable. Proton uniport in the absence of any drug-like substrate provides an additional measurement for normalization between sensors. As with methyl viologen, the magnitude of proton uniport in the presence of inward-facing and outward-facing 10-fold proton gradients was the same (Figure 7B, bottom, light colors). We then measured the magnitude of transport in the presence of simultaneous 10-fold proton gradient and 40-fold propidium gradient (pH 6, 5 μM PP^2+^ vs pH 7, 125 nM PP^2+^: Figure 7B, bottom, dark colors). We again saw increased charge transport indicating that these gradients do drive symport of PP^2+^ and H^+^. Replicate sensors for the propidium experiments can be seen in Figure S14, A-C showing how consistent these experiments are, even across sensors.

Comparing the magnitude of the transport under symport conditions to simple proton uniport provides an estimate of transport through the different pathways. Uniport of 1 or 2 H^+^ would result in net movement of +1 or +2 across the membrane during each turnover. Symport of one +2 substrate (MV^2+^ or PP^2+^) plus one H^+^ would move +3 net charge across the membrane. In the case of MV^2+^, the normalized ratio of net charge transport under proton uniport versus symport conditions was 1.37 ± 0.34 and 1.29 ±0.17 for inward- and outward-facing gradient conditions, respectively. In the case of PP^2+^, the same net charge ratio for proton uniport versus symport were 1.85 ±0.14 and 2.24 ±0.29 for inward- and outward-facing gradients. Many factors can influence this exact ratio, which will depend on the partitioning of transport among the different possible transport pathways. In particular, the actual concentration of +2 substrate relative will impact the rate of binding, and relative rate of drug binding versus alternating access in the protonated state will affect partitioning between symport and proton uniport. To further explore the contribution of different drug concentrations on partitioning between different transport pathways, we compared the signal for proton uniport in the presence of a 2-fold proton gradient (pH 7 vs 6.7) to net transport with the same 2-fold proton gradient and increasing concentrations of PP^2+^ present equally on both sides of the membrane (Figure S14D). There is a clear increase in the total transported charge with PP^2+^ present even at low concentration (0.164 μM) compared to the no-drug condition that only allows proton uniport. This confirms that symport, which moves more net charge, occurs preferentially rather than proton uniport when drug is present, since the proton driving force is unchanged but the net charge movement increases. As the PP^2+^ concentration is further increased, the total transported charge decreases, reflecting the importance of concentration-dependent on-rates for proton and small molecule substrate and the value of these on-rates compared to the alternating access rates in determining the partitioning between different transport pathways for coupled and uncoupled transport.

## Discussion

The ability to transport substrates of different charge is primarily limited to promiscuous multidrug resistance (MDR) transporters (17) and some metal transporters that can transport metal ions of different valency (18). These transporter families are involved in processes that require substrate promiscuity as well as mechanistic complexity. In this report, we probe the ability of the model MDR transporter, EmrE, to bind substrates of different charge and how this affects simultaneous binding of small molecule substrate and proton at the primary binding site defined by E14. Previous examination of EmrE-mediated transport of +1 and +2 cations found that the calculated transport stoichiometry remained the same regardless of substrate charge (4), although this conclusion is based on experiments with only two substrates, one of each charge. This led to the assumption that EmrE could not simultaneously bind drug and proton at its active site glutamates (E14_A_ and E14_B_). Our NMR pH titrations clearly shows that EmrE can bind a +2 substrate and proton simultaneously (PP^2+^, Fig. 2 and dTPP^2+^, Fig. 6), in accord with previous work from our lab showing the ability of EmrE to bind +1 drug and proton concurrently (5). Furthermore, the site-specific nature of the NMR chemical shift perturbations and comparison with E14Q mutants demonstrate that both protons and drug-like substrates of either +1 or +2 charge are bound at the primary binding site defined by the E14 residues from each subunit of the homodimer. These results indicate that neutralization of the glutamate residues does not limit the binding capacity of the transporter. The ability of this binding site to accommodate a net positive charge suggests that +3 drug-substrates also have the potential to bind and transport of such molecules should be tested in the future. This result is perhaps not entirely unexpected given recent crystal structures that show the plasticity of the EmrE binding pocket bound to different substrates (19) and NMR studies confirm the dynamic behavior of the substrate within the binding pocket (11). Indeed the NMR data we have reported previously for +1 substrates (8) and here for +2 substrates (Fig. 8) demonstrate that EmrE accommodates different substrates with changes in both the structure and rate of alternating access.

**Figure 8:**
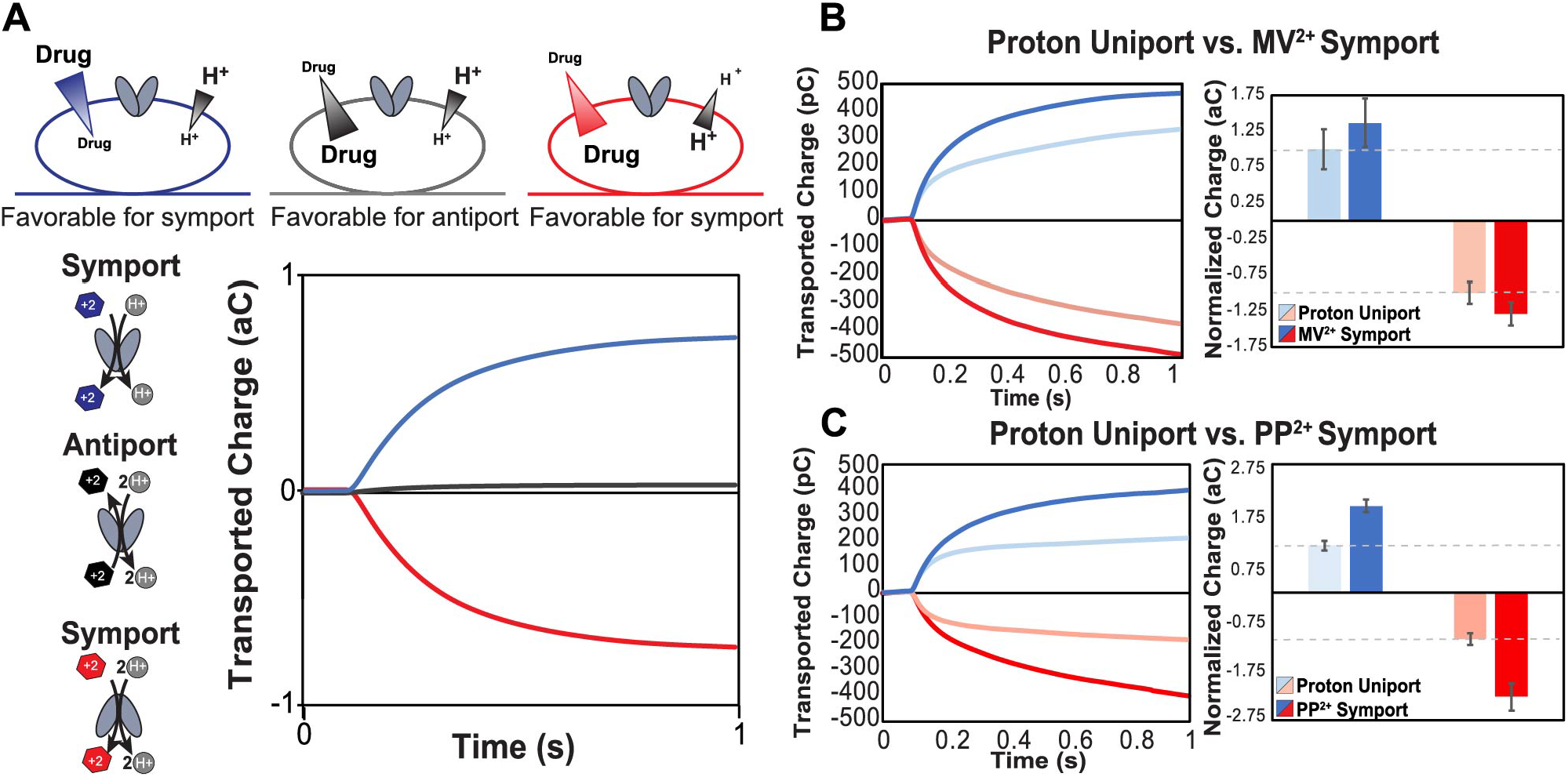
Characterizing electroneutral coupled transport of EmrE. (A) For +2 substrates, a strict 2:1 proton to drug antiport will result in net neutral transported charge with one +2 substrate moving in, while two protons move out, resulting in no signal under antiport conditions. While symport is not the most favorable transport mode for canonically antiported substrates, flux through this mode can be observed by aligning strong proton and drug gradients in the same direction. In the presence of a 10-fold proton gradient (1 pH unit), EmrE uniports protons (B, C, light color) with the same magnitude whether the gradient is inwardly directed (light blue) or outwardly directed (light red). Symport of MV^2+^ (B) or PP^2+^ (C) can be driven by simultaneous 10-fold proton and 40-fold substrate gradients and the net transport is the same magnitude but opposite direction when the gradients are inwardly directed (dark blue) or outwardly directed (dark red). Addition of the drug gradient increases net charge movement (dark vs light colors) for both MV^2+^ (B) or PP^2+^ (C), demonstrating symport of drug and proton beyond the proton uniport activity of EmrE.

Since EmrE is a loosely coupled transporter, this ability to simultaneously bind small molecule substrate and proton opens the possibility that this nominal drug/proton antiporter can also perform drug/proton symport. Our SSME experiments (Fig. 8) demonstrate that EmrE can perform symport of proton and +2 substrates, although the relative partitioning of transport through coupled and uncoupled transport pathways, and consequently the total net charge moved across the membrane, depends on the identity of the transported substrate transport substrates. These results reaffirm the complexity of EmrE activity and the need to characterize several different substrates to understand the functional behavior of a highly promiscuous and loosely coupled transporter, such as EmrE.

The ability of EmrE to bind and transport substrates with +1 or +2 is distinct from what has been reported previously for other small molecule transporters reported to transport substrates of different valency. QacA, an MDR from *S. aureus*, has been shown to confer resistance to +1 and +2 drug substrates (20, 21), but transport assays revealed non-competitive inhibition +2 drug substrates with Eth^+^, suggesting a second binding site for these compounds. The Natural resistance-associate macrophage protein (Nramp) family of metal transporters symport transition metals and protons (18), and can move metals of different charges across the membrane with different symport stoichiometries (22, 23). Structural data indicates that there are two distinct pathways in the transporter for metals and protons (24). The proton pathway does not require the transporter to undergo conformational exchange, whereas the transporter must change its conformation to transport the metal ion across the membrane. In contrast, EmrE contains a single pathway through the transporter for both +1/+2 drugs and protons and alternating access is required for transport of either or both substrates based on NMR dynamics studies in the presence and absence of drug-substrate at various pH (3, 5, 7). However, different substrates trigger different structural and dynamic changes in the transporter binding pocket (Fig. 2D, 5A, 6A), which will impact alternating access (8). This can result in flexible stoichiometries and additional complexity in EmrE’s transport mechanism, as initially proposed (5). Our SSME data in Figures 7 and 8 shows that EmrE can transport +1 or +2 drug substrates with the same stoichiometry (2 H^+^: 1 drug antiport), or with different stoichiometries (H^+^/+2 drug substrate symport) and can performed uncoupled proton uniport in addition to these different modes of coupled transport. These variable stoichiometries confirm prior results from our lab (5), and provide evidence for alternative net transport behavior by this promiscuous transporter depending on the environmental conditions. This affirms and extends the predictions of the free exchange model for EmrE transport and demonstrates the incredible breadth of promiscuity in EmrE transport, both in substrate recognition and transport mechanism.

## Experimental Procedures

### Synthesis of 1,4-phenylenebis(triphenylphosphonium) bromide (dTPP^+2^)

A 50mL heavy wall pressure vessel fitted with an internal thread of PTFE cap was charged with a magnetic stir bar, 1,4-dibromoobenzene (1.0g, 1.0 equivalent), triphenylphosphine (1.1g, 1.0 equivalent), NiCl2 hexahydrate (0.1g, 10 mol%) and ethylene glycol (20 ml). The tube was then flushed with nitrogen, then the contents were stirred at 180°C for 4 hours. The reaction mixture was cooled to room temperature, extracted with dichloromethane (3×50 mL) and the combined extracts were washed with brine solution before drying over anhydrous Na2SO4. After filtration to remove the drying agent, the solvent was removed under reduced pressure to give the crude solid product. This was recrystallized from hot water, furnishing the pure intermediate (4-bromophenyl)triphenylphosphonium bromide in a 90% yield. Next, the (4-bromophenyl)triphenylphosphonium bromide (1.8g, 1.0 equivalent), along with triphenylphosphine (0.95g, 1.0 equivalent), NiCl2 hexahydrate (0.09g, 10 mol%), and ethylene glycol (20 ml) were added under nitrogen atmosphere to a pressure vessel as before, and the reaction mixture heated and stirred at 180°C for 8 hours. Upon cooling, the reaction mixture was extracted with dichloromethane (3×50 ml) and washed with brine. After drying over Na2SO4 and filtering to remove the drying agent, the solvent was removed under reduced pressure, leaving the final product (1.76g, 64% yield). Deuterated dTPPBr2 was synthesized using the same method but using commercially available perdeuterated 1,4-dibromoobenzene and triphenylphosphine.

### Synthesis of triphenyl[4-(triphenylmethyl)phenyl]phosphonium iodide (dTPP^+^)

In a 100mL round bottom flask containing a magnetic stir bar, 4-(triphenylmethyl) aniline (2.0g, 1.0 equivalent) was dissolved in 70 mL acetone and the mixture cooled to 0°C. Concentrated HCl (7 mL) was diluted with 10 mL H_2_O, then added to the foregoing reaction mixture. After 10 m stirring at 0°C, NaNO_2_ (0.66g, 1.6 equivalents) dissolved in 5.0 mL H_2_O was added dropwise, and the mixture was stirred for 30 min at 0°C. Then, KI (1.7g, 1.62 equivalents) in 7 mL of H_2_O was added dropwise while stirring for 1h at 0°C; the solution was then allowed to warm to room temperature for over an hour. After warming to room temperature, the solution was heated for 2h at 60°C. The reaction mixture was then quenched with NaHSO_4_ (1.0g) and extracted into dichloromethane. The latter phase was separated, and the solvent was removed in vacuo, leaving 1-iodo-4-(triphenylmethyl)benzene as a tan-to-light-yellow powder. This was used without further purification in the next step. In the next step, a 50 ml heavy wall pressure vessel having a threaded PTFE cap was charged with a stir bar, 1-iodo-4-(triphenylmethyl)benzene (1.5g, 1.0 equivalent), triphenylphosphine (0.88g, 1.0 equivalent), NiCl_2_ hexahydrate (0.08g, 10 mol%), and ethylene glycol (20 ml). The tube was flushed with nitrogen, sealed, and heated with stirring for 8 hours at 180°C. The mixture was cooled to room temperature before extracting with dichloromethane (3×50 ml), the combined extracts then being washed with brine to remove any ethylene glycol which had carried over during the extraction. The dichloromethane was removed under reduced pressure, leaving 2.2 g (92% yield) of the desired product as a slightly off-white solid.

### Chemical analysis of substrates

To verify the structure of dTPP^2+^ and dTPP^+^, solution NMR spectroscopy was performed. Substrates were dissolved in either CDCl3 or DMSO-d6. ^1^H 1D and ^13^C 1D spectra were collected. SI Fig. 1 and SI Fig. 2 show representative solution NMR data collected on dTPP^2+^ and dTPP^+^, respectively, and are annotated with tentative chemical shift assignments. Data were collected using Topspin NMR 3.5 on a 600 MHz Bruker Avance III HD equipped with a 1.7 mm cryoprobe TXI (^1^H/^13^C/^15^N). For extinction coefficient determination, three samples each of dTPP^2+^ and dTPP^+^ were massed out using an analytical balance. dTPP^2+^ samples were dissolved and diluted in the water while dTPP^+^samples were dissolved in a minimal amount of DMSO before dilution in either water of 10 mM n-Dodecyl-B-D-Maltoside (DDM). Dilutions were done using a combination of Hamilton syringes and Wiretrol^®^ II pipettes (Drummond Scientific Company, Broomall, PA). UV-Vis measurements were conducted on the Evolution 201 spectrophotometer (Thermo Fisher Scientific, Waltham, MA).

### EmrE expression, purification, and reconstitution

EmrE was recombinantly expressed and purified as previously reported (8, 25). For natural abundance samples used in ITC experiments, natural abundance M9 media was used. ^2^H,^15^N-labeled WT EmrE was used in the NMR pH titration experiments, ^2^H_2_O M9 media containing 1 g/L ^15^NH_4_Cl and 0.5 g/L ^2^H/^15^N-labeled ISOGRO (Millipore-Sigma, Burlington, MA). In both cases, cultures were grown at 37 °C, shaking at 200 rpm. Protein expression was induced with 0.33 mM isopropyl β-D-1-thiogalactopyranoside (IPTG) at an OD_600_ of approximately 0.9, shaking at 200 rpm at 17 °C for 19 h.

^13^C-ILV-methyl WT EmrE was expressed in ^2^H_2_O M9 media containing 1 g/L ^15^NH_4_Cl, 2.5 g/L ^2^H-glucose, and 0.5 g/L ^2^H-labeled ISOGRO (Millipore-Sigma, Burlington, MA). The expression was conducted in the same manner as the other samples, except that approximately one hour before induction, the ILV precursors were added directly into the media. The precursors used were 120 mg/L sodium α-ketoisovalerate (^2^H-3methyl, ^13^C-4methyl; Millipore Sigma, Burlington, MA) and 70 mg/L sodium α-ketobutyrate (^2^H-3, ^13^C-methyl; Cambridge Isotope Laboratories, Tewksbury, MA). EmrE was purified also as previously reported (8, 25). Additionally, a PD-10 desalting column (GE Healthcare, Chicago, IL) was used to remove excess imidazole and buffer exchange in between the Ni affinity purification and thrombin cleavage. Thrombin cleavage was performed at room temperature overnight in buffer containing 100 mM bicine, 200-250 mM NaCl, 10 mM n-decyl β-D-maltoside, pH 8.0. The cleaved material was then purified by Superdex S-200 FPLC gel filtration as per the previous protocol. All samples were reconstituted into isotropic bicelles made of a 3:1 molar ratio of 1,2-dihexanoyl-sn-glycero-3-phosphocholine (DHPC) to 1,2-dimyristoyl-sn-glycero-3-phosphocholine (DMPC). The lipid to protein monomer ratio was 75:1. For ^13^C-methyl-ILV labeled samples, ^2^H-54-DMPC and ^2^H-22-DHPC (Avanti Polar Lipids, Alabaster, AL) were used to minimize the lipid background in the methyl region of the NMR spectra.

### Isothermal Titration Calorimetry

Isothermal titration calorimetry experiments were performed on a TA Instruments Low Volume Nano ITC instrument (TA Instruments, New Castle, DE). Data was collected using the ITCRun software at 45 °C, 350 rpm stirring. Data were fit using NanoAnalyze Software. EmrE was used as a titrand, and the ligands were used as the titrant. The monomer concentration of EmrE ranged from approximately 0.3 mM to 0.4 mM and the ligand concentrations ranged between 0.95 mM and 1.4 mM. Titrant solutions were made with a solution of bicelles with matched concentrations of DMPC and DHPC as the titrand solution. All experiments were conducted in 100 mM MOPS, 20 mM NaCl, and pH 7.0.

### NMR Spectrometry – pH titration

NMR samples for pH titrations contained ^2^H,^15^N-EmrE at 0.5-1 mM monomer concentration with an excess ligand. For titration with dTPP^2+^, an excess amount of the ligand (at least 4x monomer concentration of EmrE) was added directly to the NMR sample. For titrations with dTPP^+^, NMR samples were incubated at 45 °C with excess amounts of solid ligand overnight to ensure saturation of the binding site with the hydrophobic substrate. All samples were in 20 mM sodium acetate, 50 mM MES, 50 mM MOPS, 100 mM Bicine, 10% D_2_O, 2 mM tris(2-carboxyethyl)phosphine (TCEP), 0.01% sodium azide, and the pH was adjusted at 45 °C. Data were collected on a 900 MHz Bruker Avance III HD equipped with a 5 mm triple resonance cryoprobe (^1^H/^13^C/^15^N) using Topspin NMR 3.5 with the variable temperature (VT) set to 45 °C. Data were processed in NMRPipe (26) and analyzed in NMRFAM-Sparky (27). Amide chemical shifts were determined at each pH for 7 residues from the core of the protein. The chemical shifts were then globally fit against pH values using IgorPro (WaveMetrics, Inc, Portland, OR) to determine the pKa of EmrE bound to dTPP^2+^. The pKa of EmrE bound to dTPP^+^ could not be determined because dTPP^+^ does not bind at low pH.

### NMR Spectrometry – alternating access rate measurement

Samples used for ^13^C-edited TROSY-select ZZ exchange experiments (28) were prepared in a similar way to the pH titration samples. Data were collected on an 800 MHz Varian VNMRS DD equipped with a 5 mm cold probe (^1^H/^13^C/^15^N) using VnmrJ 4.0 with the VT setpoint at 45 °C. Spectra were collected at the following mixing times: 50 ms, 80 ms, 100 ms, 120 ms, and 150 ms for both dTPP^2+^ samples and 40 ms, 50 ms, 80 ms, 100 ms, 120 ms, and 150 ms for the dTPP^+^ sample. Data were processed in NMRPipe (26) and analyzed in NMRFAM-Sparky (27). Peak intensities were used to calculate the composite peak ratio (Ξ) using the following equation (29):

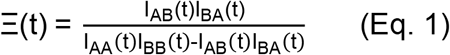

The I value corresponds to the intensity of the cross peak (AB and BA) or auto peak (AA and BB) and *t* is the mixing time of the experiment. The composite ratios were then plotted against *t* + *t*_0_, where *t* is the mixing time and *t*_0_ is the back-transfer time for these experiments, which was 9.3 ms. The alternating access exchange rate, *k_conf_*, was then determined by globally fitting the data using the following equation (5):

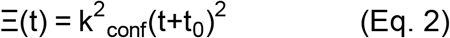

Data were fit in IgorPro (WaveMetrics Inc, Portland, OR). Errors were determined with Jackknife error analysis.

### Solid-supported membrane electrophysiology assays

WT-EmrE was expressed and purified as described with size exclusion being performed in assay buffer (50 mM MES, 50 mM MOPS, 50 mM bicine, 100 mM NaCl, 2 mM MgCl_2_, 10 mM DM, pH 7). NaOH was added to all assay buffers slowly to attain the proper pH while ensuring consistent Cl^-^ concentrations across buffer conditions. Protein was reconstituted into proteoliposomes to a 1:400 lipid-to-protein ratio in 1-palmitoyl-2-oleoyl-sn-glycero-3-phosphocholine (POPC) in pH 7 buffer, while empty liposomes underwent a simulated reconstitution. In brief, 15 mg/ml stocks of POPC were diluted in assay buffer and incubated at room temperature for 1 hour. Lipids were bath sonicated for 1 min then octyl glucoside (OG) was added to a final concentration of 0.5% w/v. Lipids were again sonicated for 30 seconds and allowed to incubate for 15 minutes at room temperature. Protein was added to the lipid solution and incubated at RT for 25 minutes, then the remaining detergent was removed with Amberlite XAD-2 as previously described (25). Uniform liposomes were obtained by extruding through a 0.2 μM membrane at least 11 times with an Avanti MiniExtruder.

Immediately before experiments, thawed liposome aliquots were diluted 2- or 4-fold, as determined for each batch by a standard assay, in pH 7 assay buffer and briefly sonicated. 10 μL of liposomes were used to prepare 3 mm sensors as previously described (14). At least five sensors each of WT and Empty liposomes were prepared and sensor capacitance and conductance values were used to select three sensors of each consistent with consistently high quality. Sensors were rinsed with at least 500 μL of internal buffer before each measurement to set the internal buffer, pH, and drug concentrations, and rinses were recorded and evaluated as described in (14). For buffer changes in the absence of drug, three washes were performed, and sensors were allowed to equilibrate in buffer for one hour before measurements. Data acquisition occurred in three stages. First, sensors were perfused with an internal buffer, then transport was initiated by perfusion of the external buffer, and finally, perfusion of the internal buffer re-equilibrated the sensors. The transported charge was obtained by integrating the current during perfusion of the external buffer, with the final 100 ms of the initial internal buffer perfusion used as the baseline. Reported data are average values of at least three sensors.

**Table I:** Summary of biophysical characterizations of dTPP derivatives.

| **Substrate** | **$K_D^{\text{pH}7}$ ($\mu\text{M}$)** | **$\Delta H^{\text{pH} 7.0}$**<br>**(kJ/mol)** | **$n^{\text{pH} 7.0}$** | **$k_{\text{conf}}$ ($\text{s}^{-1}$)** | **Source** |
| --- | --- | --- | --- | --- | --- |
| dTPP<sup>+</sup> | 8.5 ± 1.2 | -23 ± 1 | 0.54 ± 0.01 | 3.1 ± 0.1<sup>a</sup> | This paper |
| dTPP<sup>2+</sup> | 9.2 ± 0.8 | -53 ± 1 | 0.54 ± 0.01 | 3.5 ± 0.1<sup>a</sup> | This paper |
| propidium<sup>2+</sup> | 90 ± 10 | -16.3 ± 0.7 | 0.58 ± 0.06 | 13 ± 3<sup>b</sup> | (8) |

<sup>a</sup> pH 8.0
<sup>b</sup> pH 7.0

## Data Availability

Raw NMR data is deposited at the BMRB in the BMRbig database, accession number bmrbig70.

Raw biophysical and SSME data can be found at 10.17632/gs238vhsx4.1.

Until publication, use this link:

https://data.mendeley.com/datasets/gs238vhsx4/draft?a=40f2fb3a-3bae-4755-b16b-2f285f665666

## Supporting Information

This article contains supporting information.

## Acknowledgments

The authors would like to thank Dr. Darrell McCaslin and Dr. Dan Stevens of the Biophysics Instrumentation Facility at UW-Madison for the use of the Nano ITC instrument, and Dr. Marco Tonelli of NMRFAM (UW-Madison) for assisting with NMR data collection. We would also like to thank Prof. Geoff Chang for providing us with the plasmid to express EmrE. Finally, we would like to thank Dr. Nathan Thomas for his pioneering work in transport mode characterization with SSME. This study made use of the National Magnetic Resonance Facility at Madison, which is supported by NIH grant R24GM141526 (NIGMS).

## Author Contributions

Conceptualization: KAHW, JHD; Data curation: PJS, MB; Formal Analysis: PJS, MB, GSH, KAHW; Funding Acquisition: KAHW; Investigation: PJS, MB, GSH, MS; Methodology: MB (SSME), MS (chemical synthesis of TPP^+^ derivatives); Project Administration: PJS, KAHW; Resources: JHD, KAHW; Supervision: KAHW; Validation: PJS, MB, GSH, KAHW; Visualization: PJS, MB, GSH; Writing – original draft: PJS, MB; Writing – review and editing: PJS, MB, GSH, MS, JHD, KAHW

