## Supplemental Information for "Charge neutralization of the active site glutamates does not limit substrate binding and transport by EmrE"

**Supporting Information:**

Raw NMR data can be found on the BMRbig database under the accession number, bmrbig70.

Online datasets can be found on MendeleyData at:
https://doi.org/10.17632/gs238vhsx4.1

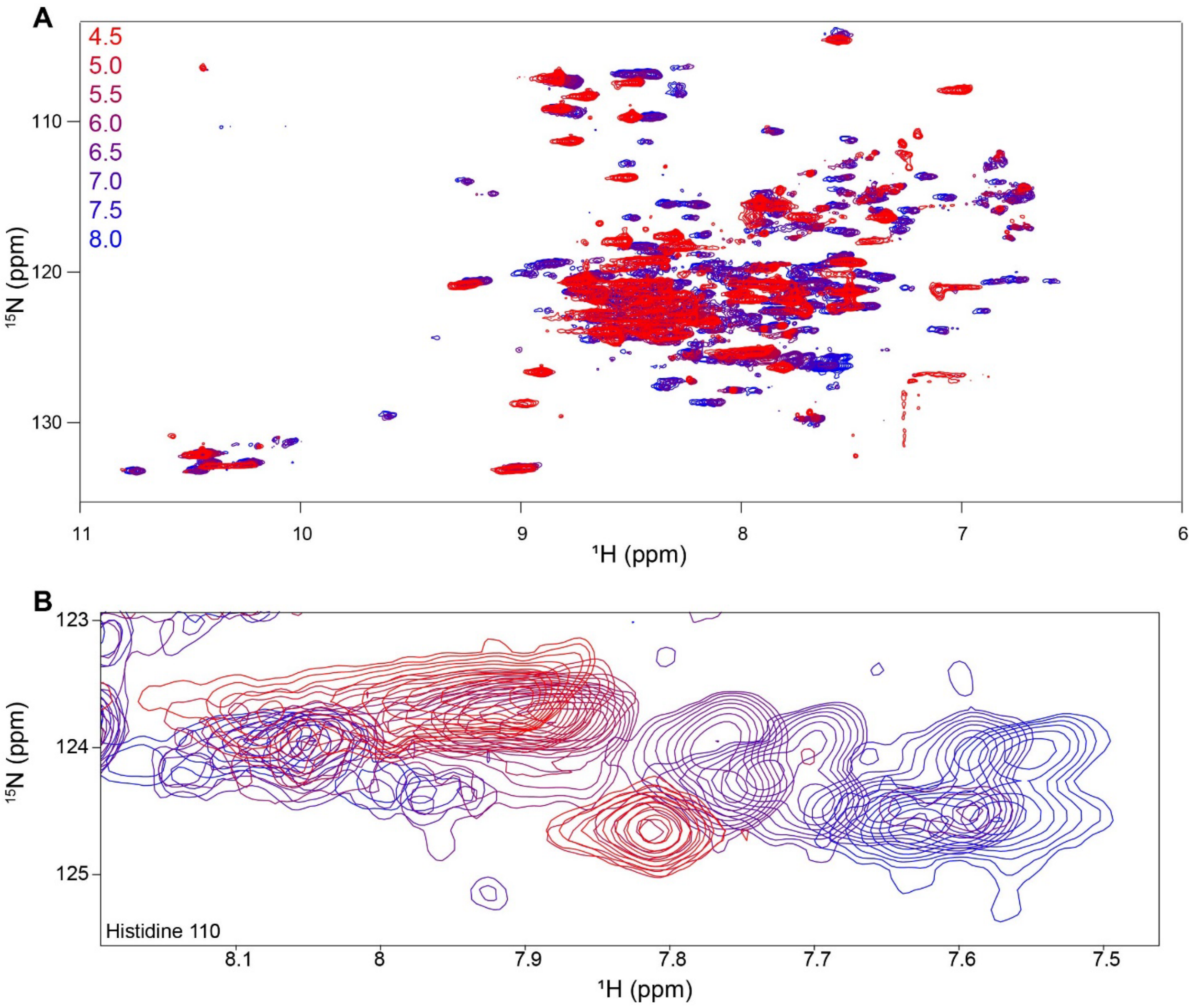


**Figure S1: pH titration of propidium-bound WT-EmrE.** A) $^{1}H^{15}N$-TROSY HSQC spectra of $PP^{2+}$-bound WT-EmrE in q=0.33 DMPC/DHPC isotropic bicelles were recorded on a Varian 800 MHz spectrometer at 45°C from low to high pH (4.5-8.0, colors as shown). These spectra exhibit chemical shift changes indicative of protonation in addition to the already-bound $PP^{2+}$. B) The shifts of H110 reveal that EmrE titrates throughout the pH range in this experiment and the lack of signal from residues, such as A10, near neutral pH are the result of line broadening from chemical exchange and not loss of protein.

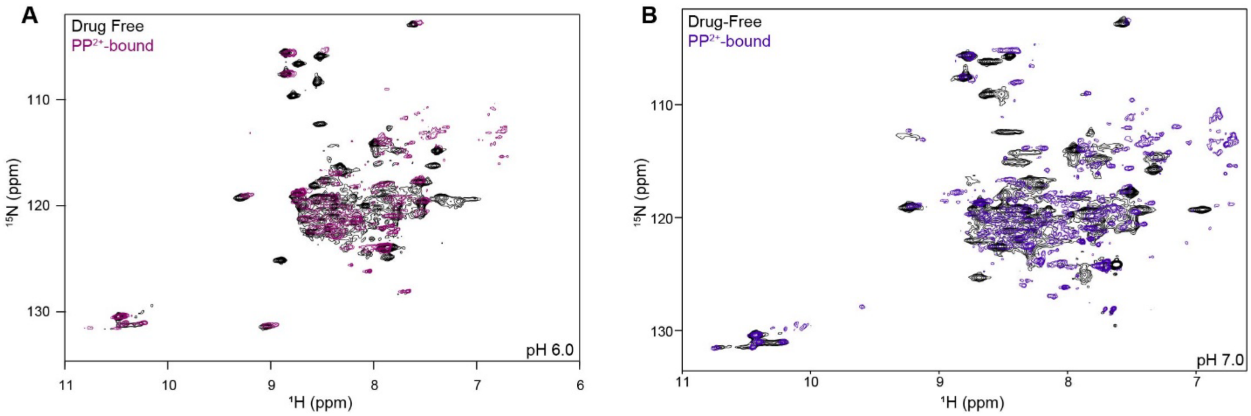


**Figure S2: Faster dynamics at physiological pH with propidium-bound WT-EmrE.** At pH 6.0 (A) and pH 7.0 (B), propidium-bound WT-EmrE has slow-intermediate timescale dynamics resulting in increased line broadening. The dynamics of the planar ligand within the transport pore combined with the exchange between different protonation states near the pKa likely contribute to this phenomenon.

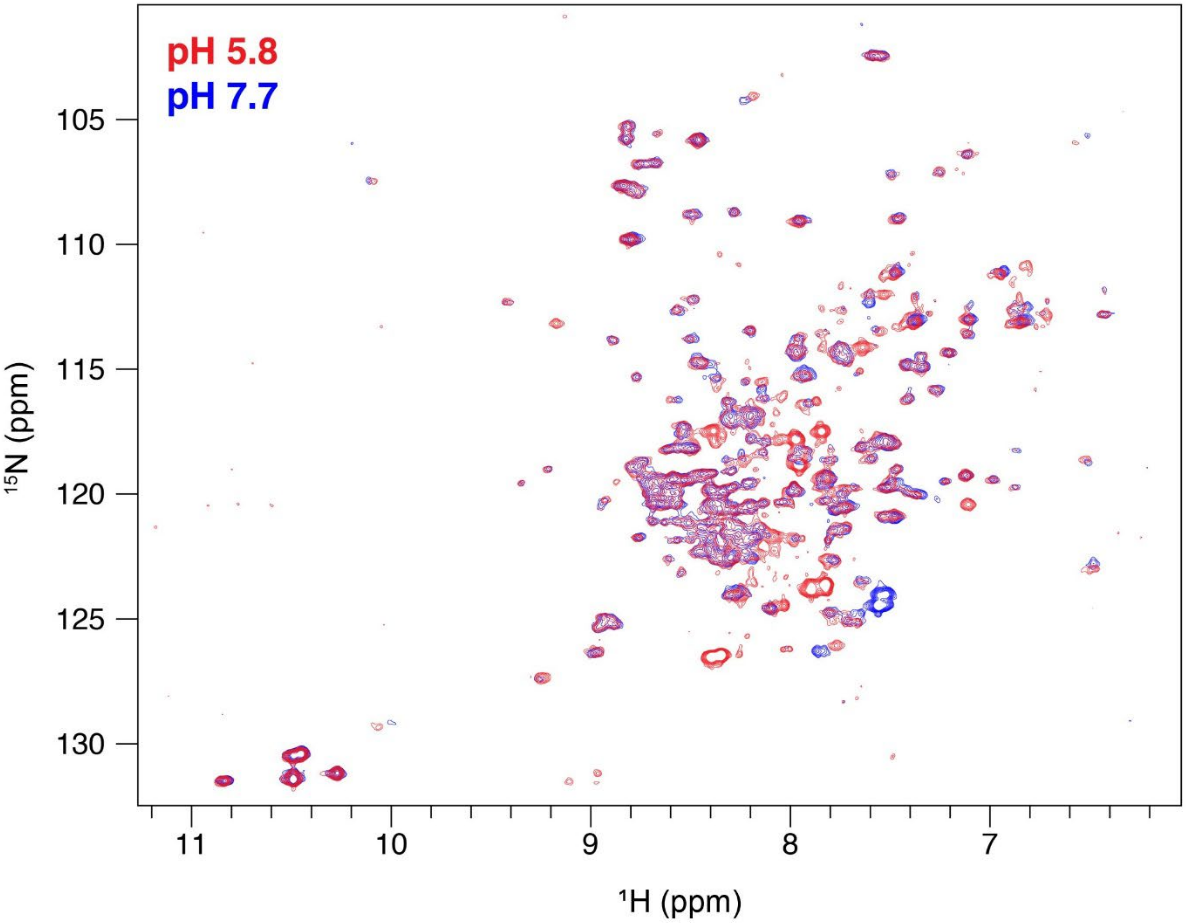


**Figure S3: E14Q-EmrE at low and high pH shows C-terminal tail titration.** E14Q-EmrE removes the effect of glutamate protonation in the active site in EmrE. Chemical shift differences between low and high pH are limited to residues in the C-terminal tail, consistent with the pKa of the C-terminal residue H110.

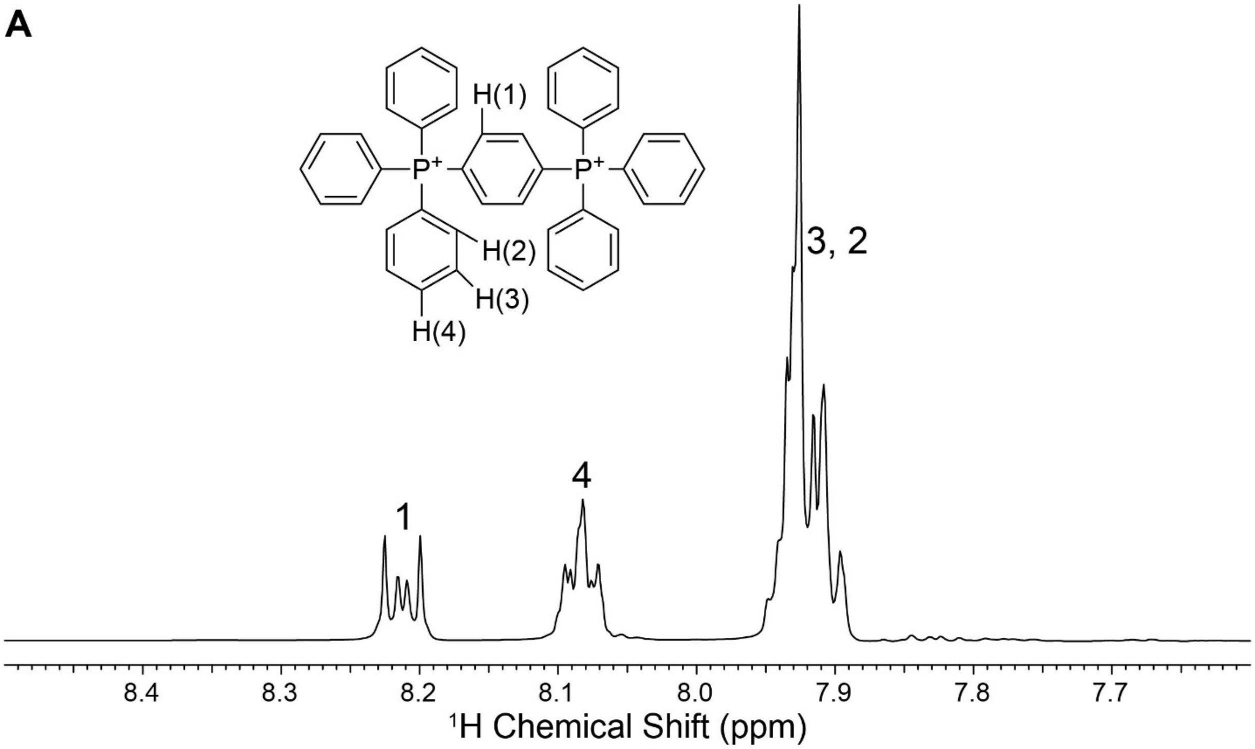


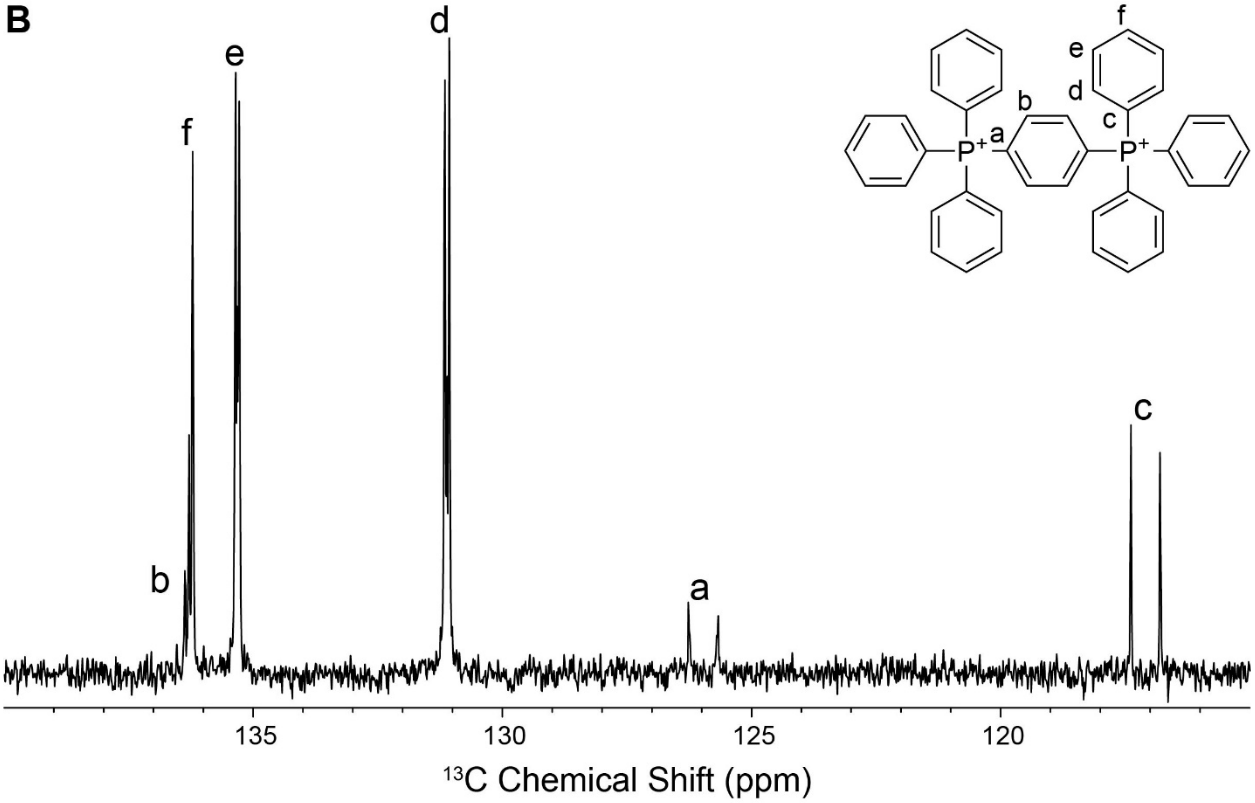


**Figure S4: 1D NMR dTPP$^{2+}$.** $^1$H (A) and $^{13}$C (B) 1D NMR spectra of dTPP$^{2+}$ confirm the symmetry of the molecule. Spectra were obtained on a Bruker Avance III 600MHz spectrometer at 25°C.

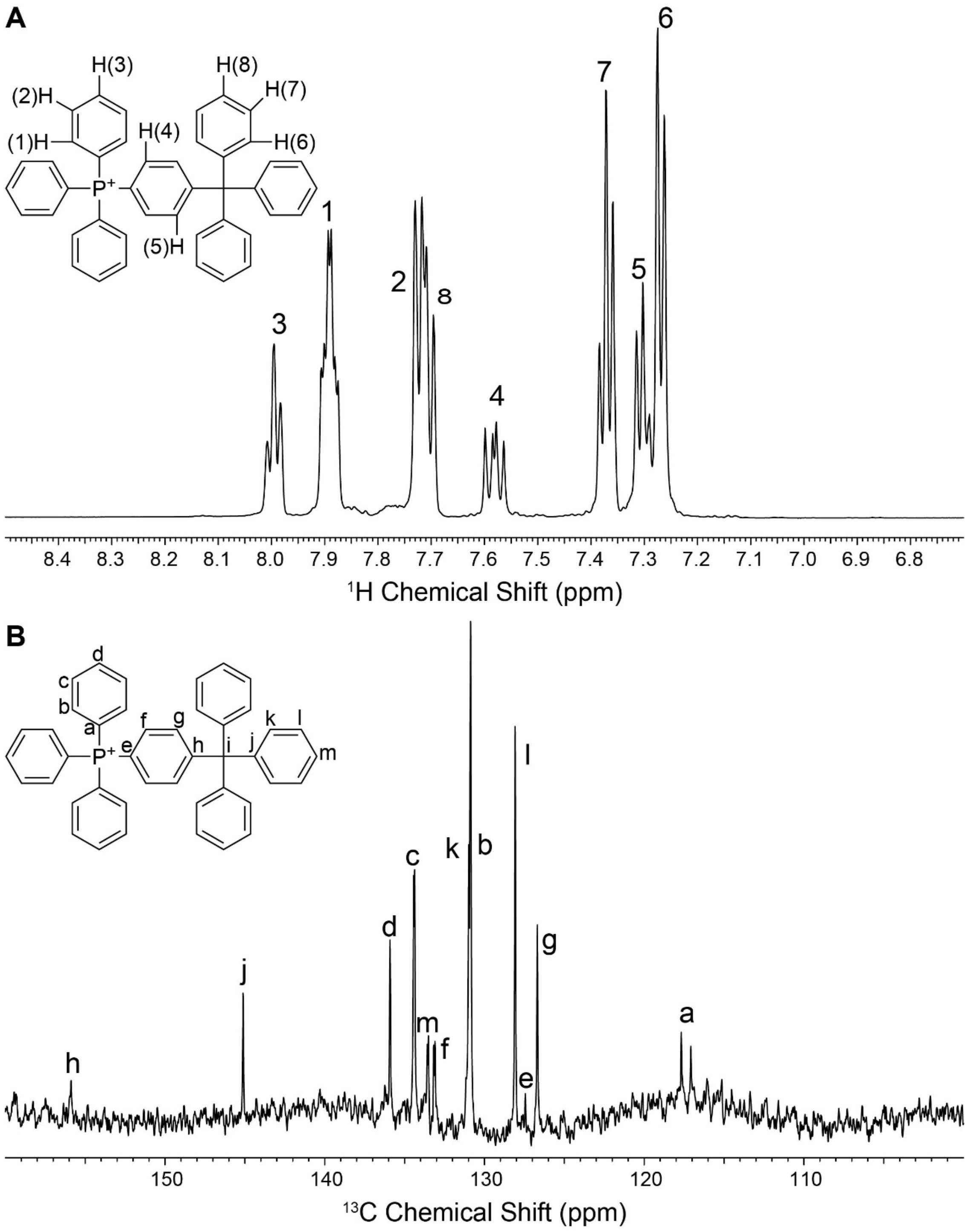


**Figure S5: 1D NMR of dTPP$^{+}$.** $^{1}$H (A) and $^{13}$C (B) 1D NMR spectra of dTPP$^{+}$ confirm the symmetry of the molecule. Spectra were obtained on a Bruker Avance III 600MHz spectrometer at 25°C.

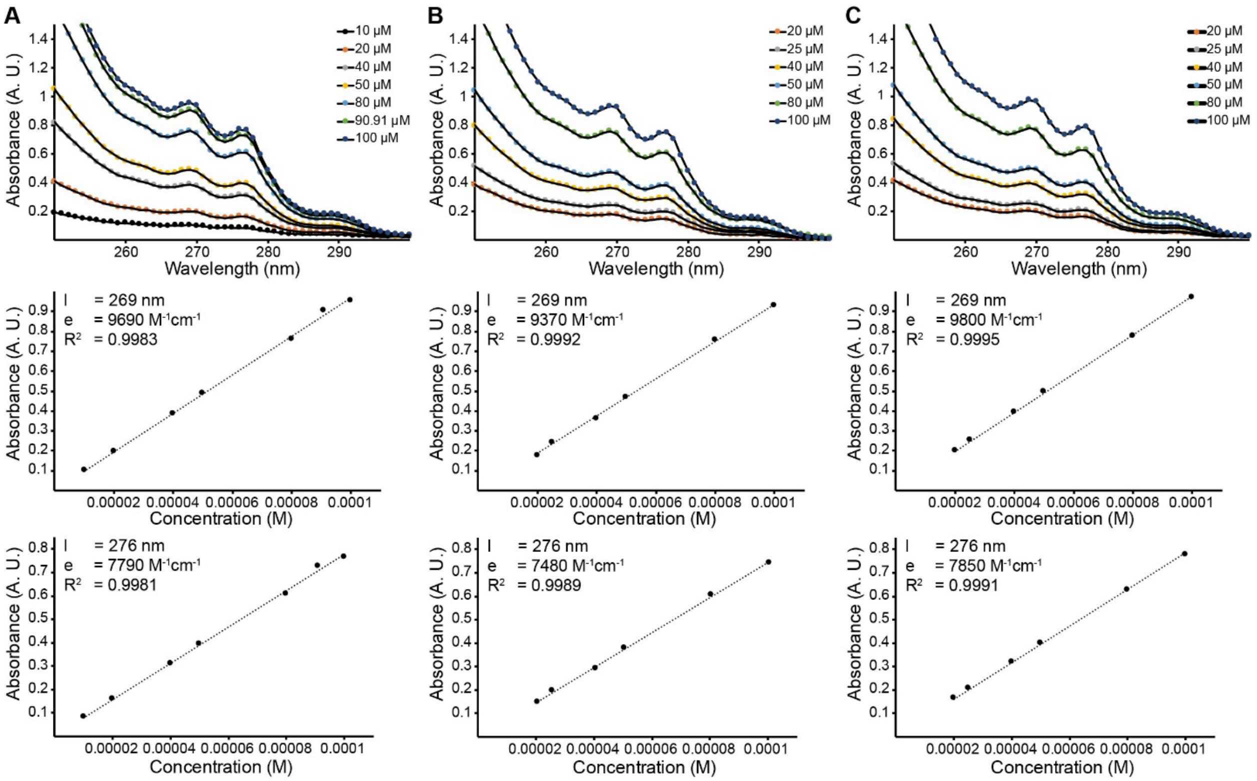


**Figure S6: Extinction coefficient trials of $dTPP^{2+}$.** Trials 1-3 (A-C, respectively) of extinction coefficient experiments for $dTPP^{2+}$. A summary of the values from this analysis can be found in Table S1.

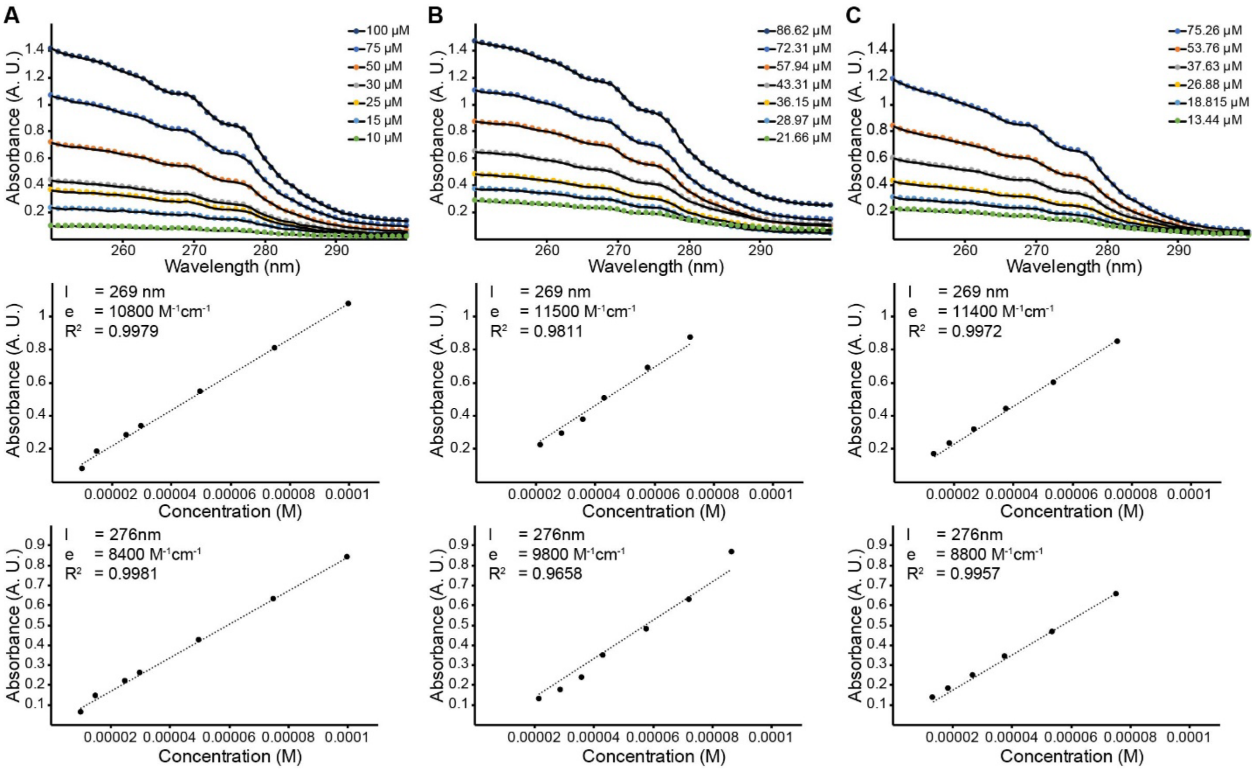


**Figure S7: Extinction coefficient trials of dTPP$^+$.** Trials 1-3 (A-C, respectively) of extinction coefficient experiments for dTPP$^+$. A summary of the values from this analysis can be found in Table S1.

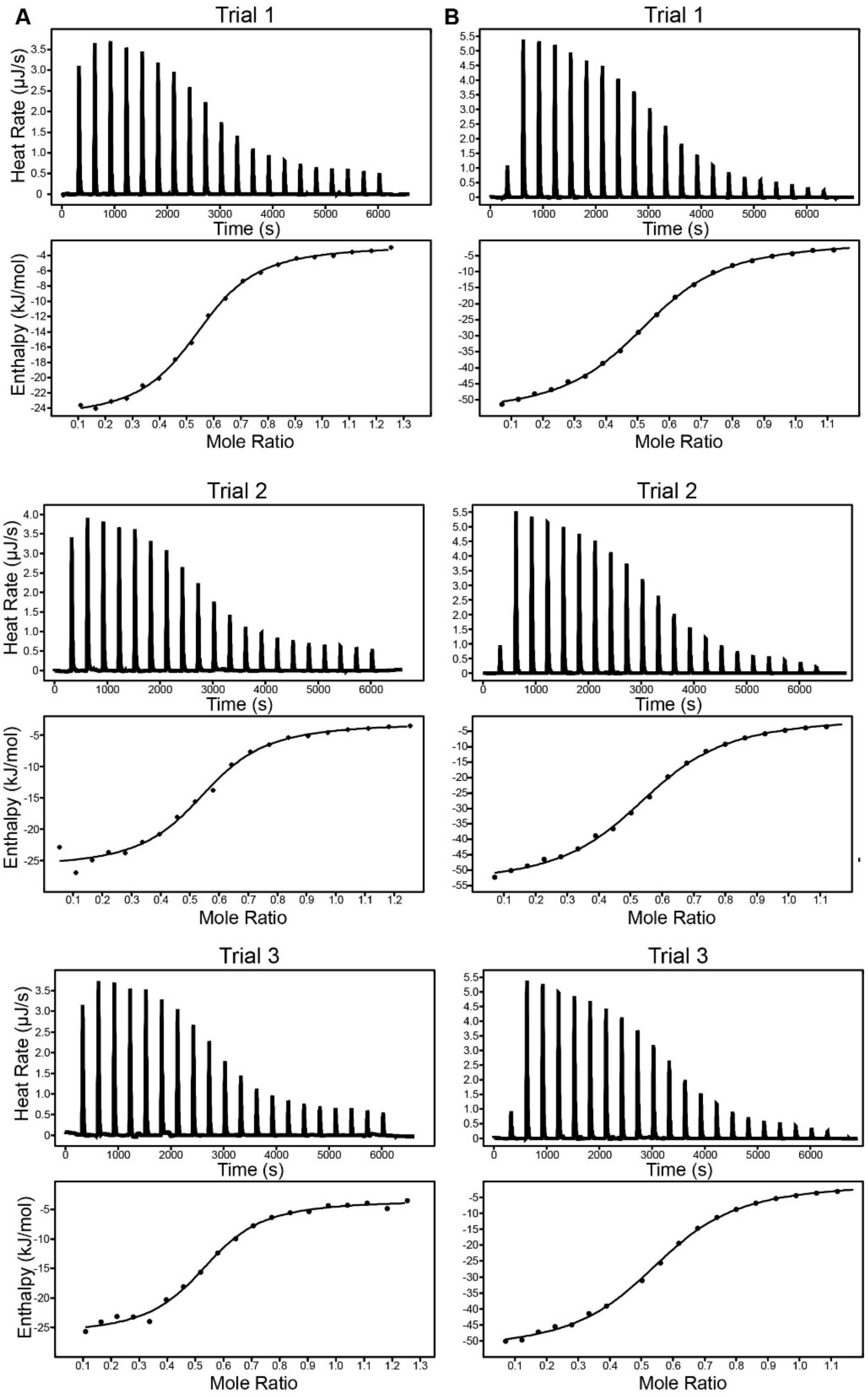


**Figure S8: ITC trials of dTPP derivatives.** ITC curves from Trials 1-3 of $dTPP^{2+}$ (A) and $dTPP^{+}$ (B) at pH 7.0.

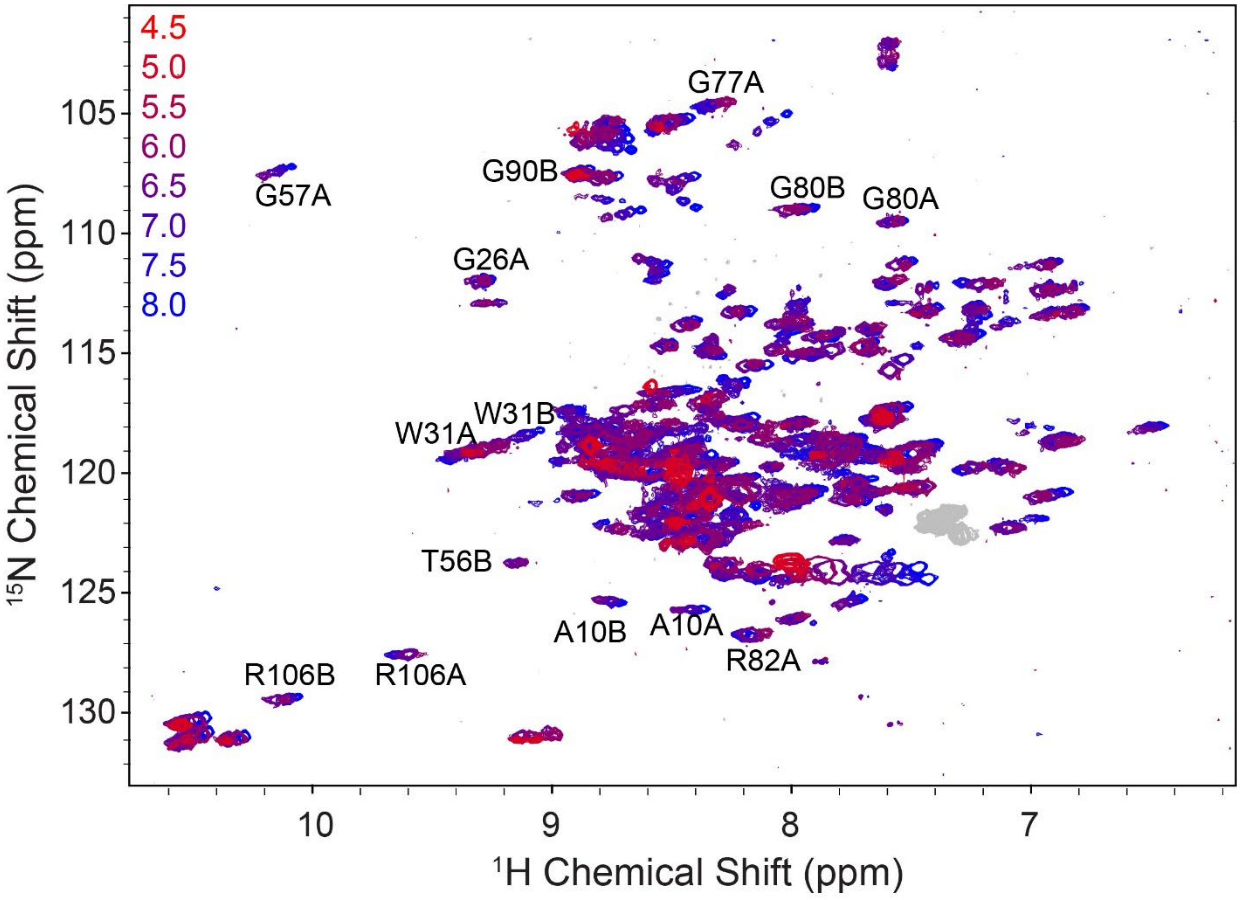


**Figure S9: pH titration of dTPP⁺-bound WT-EmrE.** $^{1}H^{15}N$-TROSY HSQC spectra of dTPP⁺-bound WT-EmrE in q=0.33 DMPC/DHPC isotropic bicelles were recorded on a Varian 800 MHz spectrometer at 45°C from low to high pH (4.5-8.0, colors as shown). These spectra exhibit chemical shift changes indicative of protonation in addition to the already bound dTPP⁺. Peaks were assigned using original assignments from TPP⁺-bound WT-EmrE pH titrations.

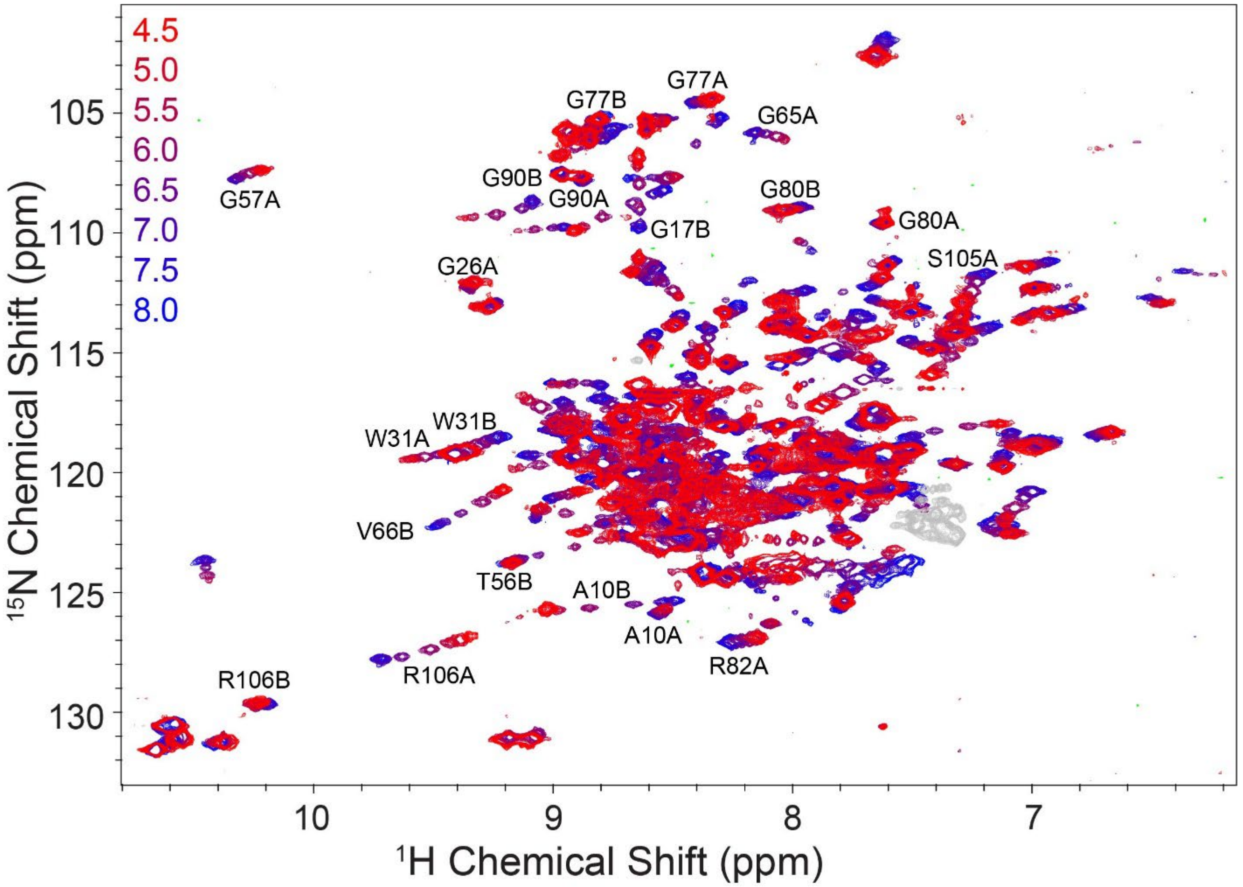


**Figure S10: pH titration of $dTPP^{2+}$-bound WT-EmrE.** $^{1}H^{15}N$-TROSY HSQC spectra of $dTPP^{2+}$-bound WT-EmrE in q=0.33 DMPC/DHPC isotropic bicelles were recorded on a Varian 800 MHz spectrometer at 45°C from low to high pH (4.5-8.0, colors as shown). These spectra exhibit chemical shift changes indicative of protonation in addition to the already bound $dTPP^{2+}$. Peaks were assigned using original assignments from $TPP^{+}$-bound WT-EmrE pH titrations.

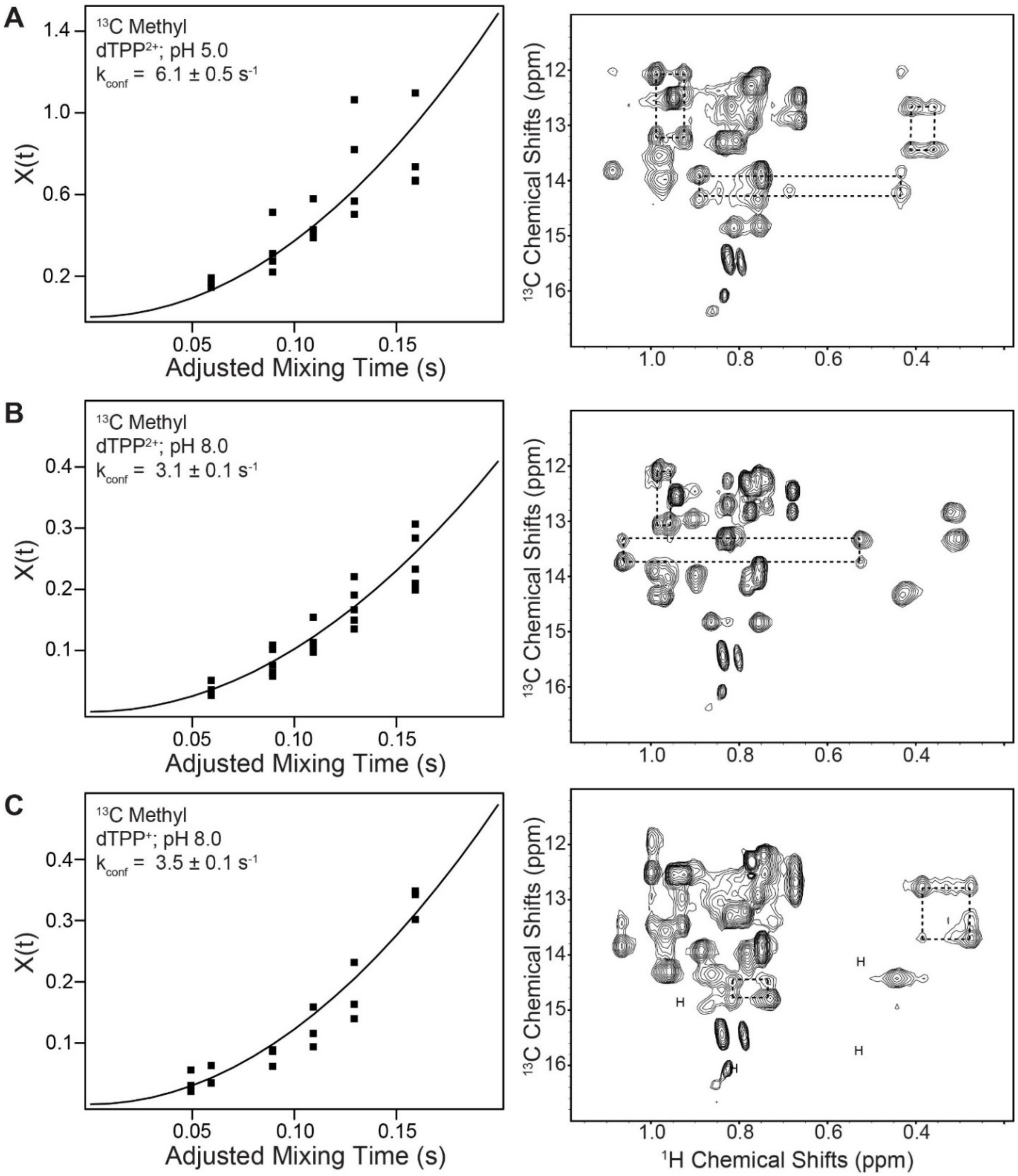


**Figure S11: Methyl ZZ-exchange spectroscopy of dTPP derivatives at low and high pH.** The rate of alternating access of ILV-labeled WT-EmrE saturated with either $dTPP^{2+}$ (pH 5.0, A; pH 8.0, B) or $dTPP^{+}$ (pH 8.0, C) were analyzed using methyl ZZ-exchange spectroscopy. Sample spectral planes with intermediate mixing times are shown on the right and boxes are drawn to highlight methyl resonances for which auto peaks for state A and state B and the cross peaks are all sufficiently resolved to use in quantitative analysis. The auto and cross peak ratio, calculated as described in the methods, is shown as a function of mixing time on the left along with the best fit.

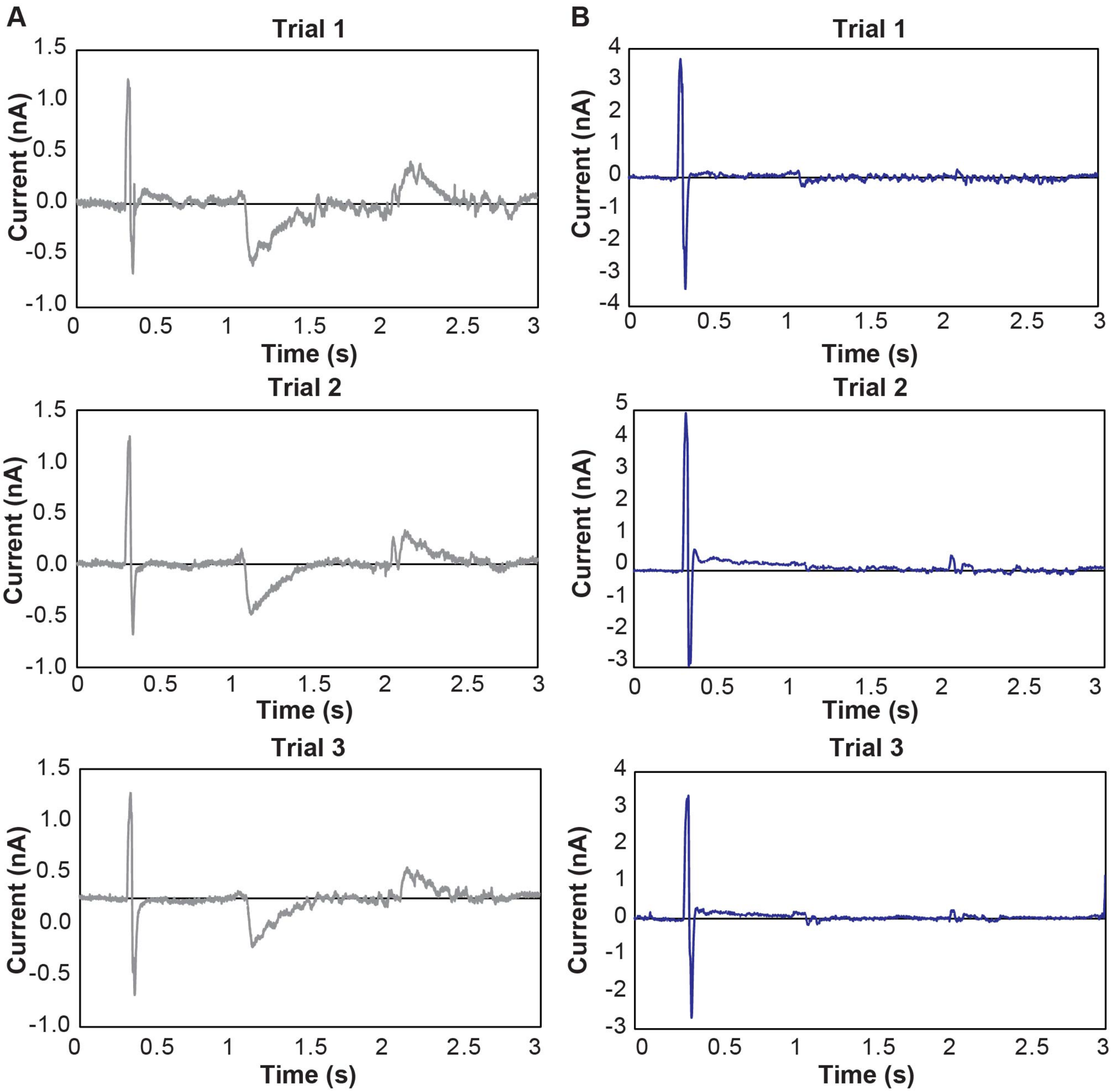


**Figure S12: SSME trials of $Eth^+$ transport.** Trials of 1 μM Ethidium transport at pH 7 using three separately prepared sensors each of WT-EmrE proteoliposomes (A) and empty liposomes (B). Transport currents for the WT sensors overlay well demonstrating reproducibility, while Empty sensors show minimal signal confirming that minimal interactions are taking place with the lipid membrane at this concentration.

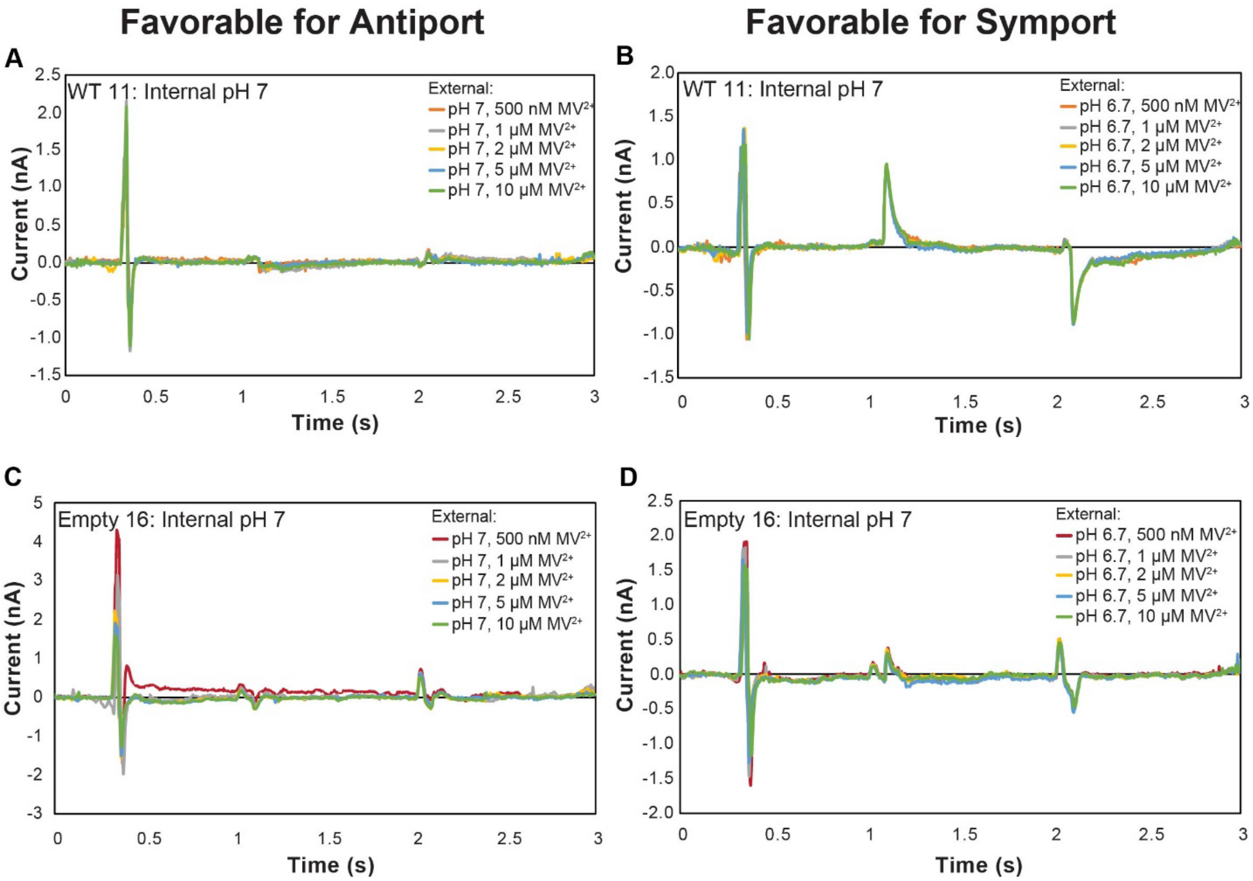


**Figure S13: SSME transport of the +2 $MV^{2+}$.** When conditions favorable for driving antiport are applied with the +2 substrate $MV^{2+}$, minimal is detected as expected for net neutral antiport of two protons out of the liposome for each $MV^{2+}$ molecule that enters the liposome (A). The same sensor was perfused with buffers with the same $MV^{2+}$ concentrations but pH 6.7 to create 2-fold proton gradients in the same direction as the drug gradient. This drives symport (B). A sensor containing empty liposomes was perfused with the same buffering conditions to confirm that the signal observed is due to EmrE-mediated transport (C, D).

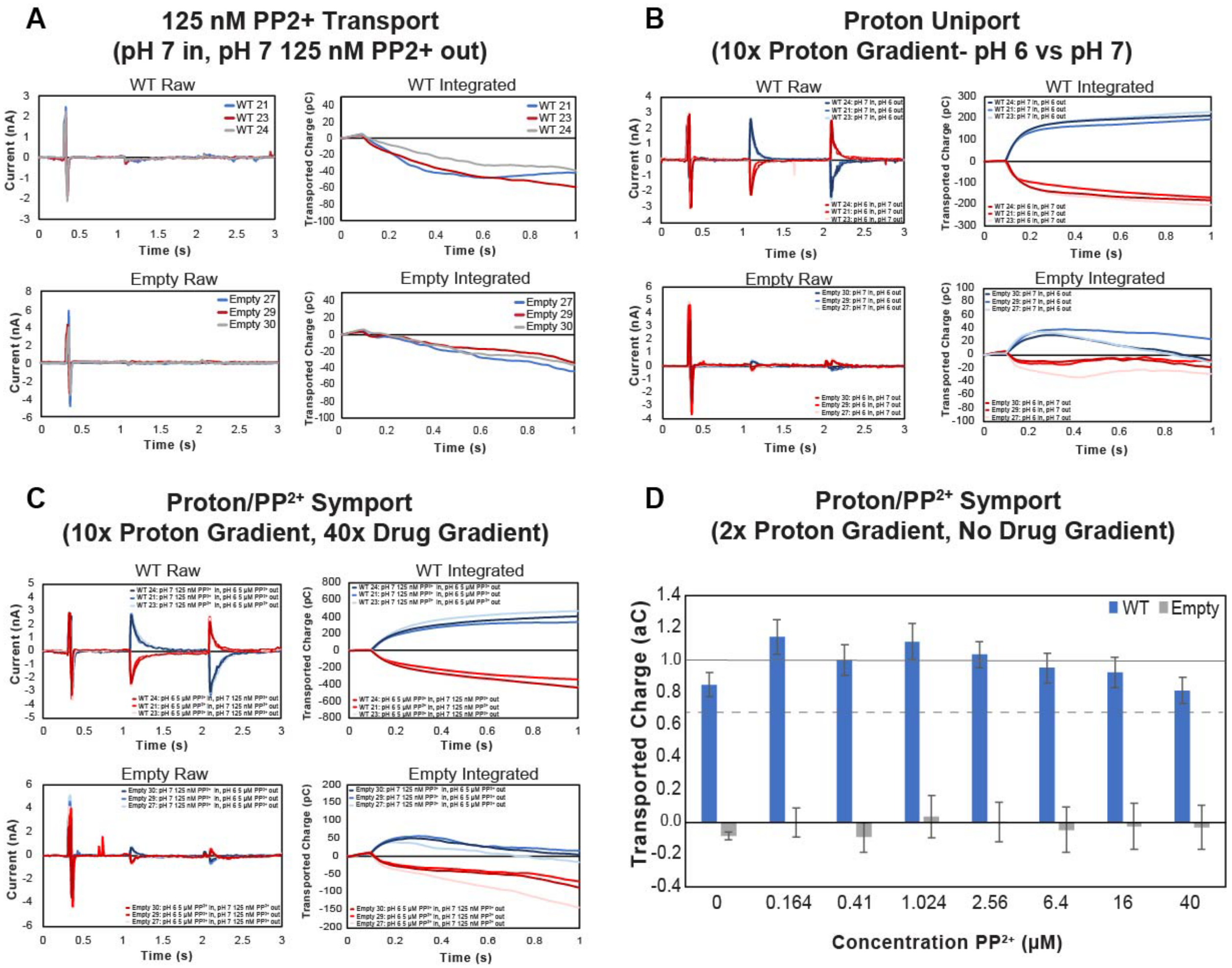


**Figure S14: SSME transport of the +2 $PP^{2+}$.** Overlays of the different sensors averaged for replicates of $PP^{2+}$ transport (A), proton uniport (B), and proton/PP2+ symport (C) show a high degree of reproducibility for the assay. Note the different axis scale in each panel. Normalized transported charge for proton/PP2+ symport under a lesser proton gradient (2x) shows how transported charge can vary drastically as the kinetics of transport are affected by substrate concentration as well as the magnitude of the gradients present (D).

**Table S1: Extinction coefficient trials**

| | **$dTPP^{2+}$ $\varepsilon_{269}$ ($M^{-1}cm^{-1}$)** | **$dTPP^{2+}$ $\varepsilon_{276}$ ($M^{-1}cm^{-1}$)** | **$dTPP^{+}$ $\varepsilon_{269}$ ($M^{-1}cm^{-1}$)** | **$dTPP^{+}$ $\varepsilon_{276}$ ($M^{-1}cm^{-1}$)** |
|---|---|---|---|---|
| Trial 1 | 9692 | 7793 | 10778 | 8434 |
| Trial 2 | 9366 | 7481 | 12188 | 9672 |
| Trial 3 | 9796 | 7852 | 11361 | 8804 |
| Average | 9618 | 7708 | 11442 | 8970 |
| Standard Deviation | 224 | 199 | 709 | 635 |
| Consensus Value | 9600 ± 200 | 7700 ± 200 | 11400 ± 700 | 9000 ± 600 |

**Table S2: ITC trial summary**

| | $K_D$ | *n* | ΔG | ΔH | ΔS | $[EmrE]_{monomer}$ | [Substrate] | *c*-value |
|---|---|---|---|---|---|---|---|---|
| | μM | | kJ/mol | kJ/mol | J/mol•K | mM | mM | |
| $dTPP^{2+}$ Trial 1 | 8.38 ± 1.38 | 0.54 ± 0.01 | -30.9 ± 0.4 | -22.5 ± 0.7 | 26.42 | 0.38 | 1.36 | 25 ± 4 |
| $dTPP^{2+}$ Trial 2 | 11.00 ± 3.55 | 0.54 ± 0.02 | -30.2 ± 0.9 | -24.8 ± 1.7 | 16.97 | 0.38 | 1.36 | 19 ± 6 |
| $dTPP^{2+}$ Trial 3 | 7.59 ± 2.91 | 0.53 ± 0.02 | -31.2 ± 1.0 | -22.7 ± 1.6 | 26.75 | 0.38 | 1.36 | 27 ± 10 |
| $dTPP^{+}$ Trial 1 | 9.14 ± 1.03 | 0.53 ± 0.01 | -30.7 ± 0.3 | -53.8 ± 1.3 | -72.5 | 0.29 | 0.96 | 17 ± 2 |
| $dTPP^{+}$ Trial 2 | 9.47 ± 1.87 | 0.54 ± 0.01 | -30.6 ± 0.5 | -54.2 ± 2.4 | -74.2 | 0.29 | 0.96 | 16 ± 3 |
| $dTPP^{+}$ Trial 3 | 8.99 ± 1.57 | 0.56 ± 0.01 | -30.7 ± 0.5 | -52.4 ± 2.0 | -68.15 | 0.29 | 0.96 | 18 ± 3 |
| $^{2}H$-$dTPP^{2+}$ Trial 1 | 13.42 ± 2.58 | 0.52 ± 0.01 | -29.7 ± 0.5 | -27.8 ± 1.3 | 6.02 | 0.40 | 1.20 | 15 ± 3 |
| $^{2}H$-$dTPP^{2+}$ Trial 2 | 12.48 ± 2.04 | 0.51 ± 0.01 | -29.9 ± 0.4 | -27.6 ± 1.0 | 7.08 | 0.40 | 1.20 | 16 ± 3 |
| $^{2}H$-$dTPP^{2+}$ Trial 3 | 12.00 ± 1.57 | 0.51 ± 0.01 | -30. 0 ± 0.3 | -27.7 ± 0.8 | 7.25 | 0.40 | 1.20 | 17 ± 2 |
